# Pan-cancer landscape of AID-related mutations, composite mutations and its potential role in the ICI response

**DOI:** 10.1101/2021.06.26.447715

**Authors:** Isaias Hernández Verdin, Kadir C. Akdemir, Daniele Ramazzotti, Giulio Caravagna, Karim Labreche, Karima Mokhtari, Khê Hoang-Xuan, Matthieu Peyre, Franck Bielle, Mehdi Touat, Ahmed Idbaih, Alex Duval, Marc Sanson, Agustí Alentorn

## Abstract

Activation-induced cytidine deaminase, *AICDA* or AID, is a driver of somatic hypermutation and class-switch recombination in immunoglobulins. In addition, this deaminase belonging to the APOBEC family, may have off-target effects genome-wide, but its effects at pan-cancer level are not well elucidated. Here, we used different pan-cancer datasets, totaling more than 50,000 samples analyzed by whole-genome, whole-exome or targeted sequencing. AID synergizes initial hotspot mutations by a second composite mutation. Analysis of 2.5 million cells, normal and oncogenic, revealed *AICDA* expression activation after oncogenic transformation and cell cycle regulation loss. AID mutational load was found to be independently associated with favorable outcome in immune-checkpoint inhibitors (ICI) treated patients across cancers after analyzing 2,000 samples. Finally, we found that AID related neoepitopes, resulting from mutations at more frequent hotspots if compared to other mutational signatures, enhance *CXCL13*/*CCR5* expression, immunogenicity and T-cell exhaustion, which may increase ICI sensitivity.

**In Brief:** A combined bulk and single cell multi-omic analysis of over 50,000 patients and 2.5 million cells across 80 tumor types reveals oncogenic acquired AICDA expression inducing composite mutations and clonal immunogenic neoepitopes that are associated with favorable outcome in patients treated by immune-checkpoint inhibitors.

**Highlights:**

• Pan-cancer analysis of AID mutations using > 50,000 samples, 2,000 ICI treated cases and 2.5 million cells with genome, exome and transcriptome data
• Oncogenic transient *AICDA* expression induces mutations mainly during transcription of its off-target genes in virtually all cancers
• AID is implicated in composite mutations on weakly functional alleles and immunogenic clonal neoepitopes at hotspots with greater positive selection
• AID mutational load predicts response and is associated with favorable outcome in ICI treated patients

---

Activation-induced cytidine deaminase (AID, encoded by *AICDA*) was initially described as the driver of somatic hypermutation (SHM), which diversifies the variable (V) domains of immunoglobulin genes (Honjo et al., 2002, 2004), and later as responsible for class-switch recombination (CSR) in activated B cells in germinal centers (Muramatsu et al., 2000). However, off-target AID activity also contributes to “aberrant” mutagenesis throughout the genome and has been mainly described in lymphomas and other hematological cancers (Pasqualucci et al., 2008; Rustad et al., 2020), but only in some solid cancers (Komori et al., 2008; Sapoznik et al., 2016; Sawai et al., 2015; Shimizu et al., 2014). More broadly, AID belongs to a large family of enzymes called apolipoprotein B mRNA editing enzyme, catalytic polypeptide-like (APOBEC) which are considered as a source of somatic mutations in a variety of cancers (Alexandrov et al., 2013; Roberts et al., 2013; Swanton et al., 2015). Nevertheless, a comprehensive characterization of the role of AID-related mutations at pan-cancer level and its potential mutational and clinical implications has not been performed.

To test this, we used a large list of pan-cancer bulk sequencing datasets including The Cancer Genome Atlas (TCGA) (Ellrott et al., 2018), the Pan-Cancer Analysis of Whole Genomes (PCAWG) (Campbell et al., 2020), MSK-IMPACT (Gorelick et al., 2020; Zehir et al., 2017), different pan-cancer pediatric projects (Gröbner et al., 2018; Ma et al., 2018), multiple hematological cancer datasets (Papaemmanuil et al., 2016; Reddy et al., 2017; Tyner et al., 2018) and we even included some studies of non-human mammals (Amin et al., 2020; Gardner et al., 2019; Wong et al., 2019a), summing up to around 49 thousand samples. We found AID somatic mutations having around a 5% frequency in virtually all cancers (human and not-human), showing stronger activity at transcriptionally active domains and synergizing initial hotspot mutations by a second composite mutation.

Interestingly, the APOBEC mutational signature, defined as single-base substitution (SBS) somatic signatures SBS2 and SBS13 according to Alexandrov (Alexandrov et al., 2020), has been proposed as a biomarker for ICI response in non-small cell lung cancer (NSCLC) but also in other cancers (Litchfield et al., 2021a; Wang et al., 2018); additionally, recent evidence suggests that APOBEC-related modification of the tumor cell immunopeptidome may increase the immunogenicity to drive anti-tumor therapy (Driscoll et al., 2020). Therefore, we used more than 2.000 ICI-treated samples (Miao et al., 2018a; Pender et al., 2021a; Samstein et al., 2019a), finding AID-related fraction of mutations as an independent prognostic value to ICI after adjusting by TMB and APOBEC signature.

Additionally, we analyzed 18 tumor types across 41 studies plus three studies covering the human, mouse and PBMC atlas of normal cells, all at single cell resolution to demonstrate that *AICDA* expression is acquired after malignant transformation (Han et al., 2018a, 2020a; Zheng et al., 2017b). Overall, we used more than 50.000 samples covering more than 80 tumor types at bulk level and close to 2.5 million cells at single cell resolution to thoroughly describe the landscape of AID-related mutations (see Figure 1A, Figure 7 and Supplementary Tables 1-2).

**Figure 1.**
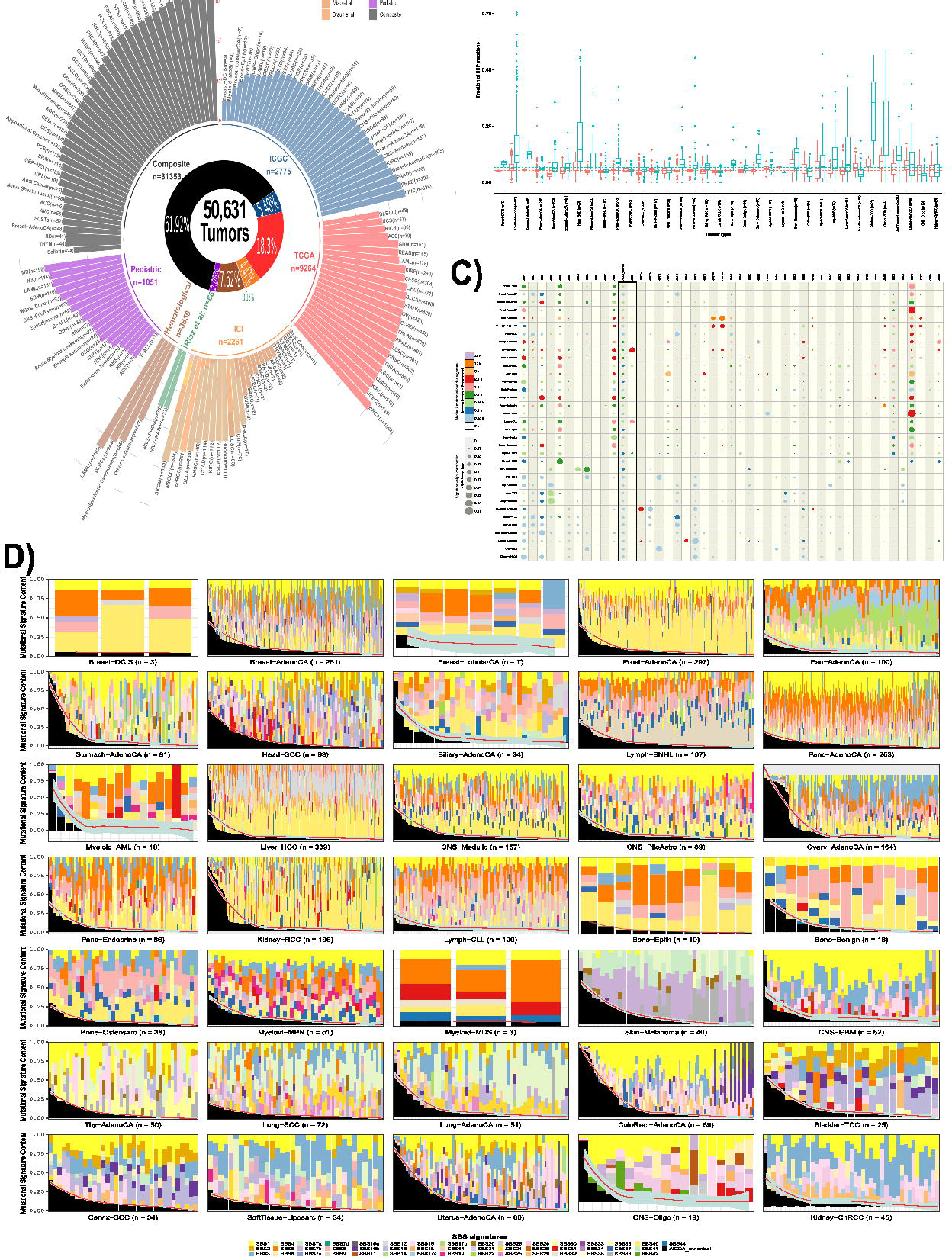
Pan-cancer landscape of AID-related mutations. Panel A, samples’ distribution of the different cohorts used in the study (51,631 tumors) where each bar is a tumor type or subgroup (only Riaz et al cohort); for the ICI cohort stacked bars represent the different studies. The complete 88 tumor types’ abbreviations are presented in Supplementary Table 2. Panel B, frequency of the fraction of mutations attributed to *AICDA* motifs or *APOBEC* motifs in the ICGC. Panel C, distribution of Single Base Substitutions (SBS) according to the COSMIC signatures in the ICGC dataset. Panel D, distribution of the SBS signatures contribution in the different cancer types of the ICGC dataset per sample where AICDA signature is represented in black with a smooth regression line in red.

## Results

### Landscape of AID-related mutations at pan-cancer level

While APOBEC signature has been found crucial in driving tumorigenesis, drug resistance and *kataegis* events in over half of human tumors, a closely related enzyme, AID, has not been covered at pan-cancer scale (Langenbucher et al., 2021; Roberts et al., 2013). We found AID-related mutations in the vast majority of cancers that we studied (Figure 1). Overall, the AID-related mutations were found in roughly 5.2% (5.1-5.3% at 95% confidence interval [CI]) and 6.6% (6.5-6.8% at 95% CI) in APOBEC mutations (Figure 1B). When we included the AID-related mutations to the SBS COSMIC signatures, we also found similar results at pan-cancer level using the WGS ICGC dataset (Figure 1C-D), TCGA, MSKCC cohort and different pediatric datasets (Supplementary Figures 1-2). Conversely, as expected, the frequency of AID-related mutations was slightly higher in hematological cancers at approximately 8% (Supplementary Figure 2F). Intriguingly, the AID-mutations were also identified in canine melanoma, glioma and osteosarcoma at a frequency of 6.0%, 4.7% and 2.9%, respectively (Supplementary Figure 2G). Additionally, in regard to the genomic distribution of *AICDA* motifs (in the normal genome) and mutations (within tumors), for the majority of tumor types the highest density of mutations were located in chromosome 5, in which *GPR98* and *DNAH5* were frequently affected, followed by chromosome 17 and 2 (Supplemental Figures 3-6). However, the number of *AICDA* motifs in the genome is higher in the latest (0.01 and 3.2x10^-4^, respectively; FDR corrected p-value Wilcoxon-test) but due to its chromosome length (Supplementary Figure 7A-B). In DLBCL most commonly affected chromosomes involved the presence of either immunoglobulin related genes: *IGH* (chr14), *IGL* (chr22), *IGK* (chr2) or genes already related with off-target AID activity: *PIM1*, *IRF4*, *HIST1H1C* (chr6; Supplementary Figure 7C-F) (Lossos et al., 2004). Interestingly, within the driver genes context hematological cancers (i.e. Lymph-BNHL, DLBCL) and medulloblastoma had the highest signature contribution of AICDA provoked mutations. Furthermore, among the involved targets, *TP53,* in all cohorts; *IDH1,* in hematological cancers, GBM and LGG; and *PIK3* genes (TCGA and ICGC cohorts), were recurrently altered (Supplementary Figure 8). These results were also confirmed using a selection intensity approach of every somatic in the ICGC dataset, showing a higher selection intensity of *PIK3CA*, *NFE2L2* but also in “minor” *IDH1* mutations (i.e. not R132H) and *PTEN* (Supplementary Figure 9).

*AICDA* expression and AID-related mutations were not correlated, and only in THCA were slightly positively correlated (Rho=0.18, p_adj_=0.01), suggesting that *AICDA* is not constitutively activated in any cancer . The AID signature was more frequently negatively correlated with the tumor mutation burden (TMB) of cancers from TCGA (i.e. in ACC, KIRP, KIRC, LIHC, LUAD, OV and THCA). AID mutations were found in younger patients when compared to APOBEC mutations (median age 61 years vs 65 years p = 0.009). However, within AID mutations, we did not find any difference of age according to gender, but only in the APOBEC related mutations where these mutations were found in elderly men compared to women (p = 7.3x10^-7^, Wilcoxon-test, Supplementary Figure 10). In brief, we found AID activity leaves important DNA footprints across human and not-human tumors, including driver genes.

### Relationship of AID-related mutations with immune related signatures and the presence of viral genome

Taking together that normal *APOBEC* expression is induced after viral exposition, recent findings of oncogenic *APOBEC* expression correlating with positive HPV infection in HNSC (Cannataro et al., 2019) and studies showing correlation between AID-related mutations with chronic infections (i.e. not viral), like with *Helicobacter pylori* (*H. pylori*) in precancerous stages of stomach cancer or Plasmodium infection in B cell lymphoma (Robbiani et al., 2015; Shimizu et al., 2014), we sought to analyze the potential relationships between the presence of AID-related mutations, the *AICDA* expression and the presence of different oncogenic viruses at pan-cancer level using the TCGA dataset (Figure 2A-C). In addition, we also analyzed their relationship with different immune-related cells, obtained by deconvolution. The AID-related mutations only showed a significant enrichment in STAD tumors with negative Epstein-Barr virus (EBV) infection (Figure 2C). Intriguingly, *AICDA* expression was significantly higher in BLCA, HNSC and CESC with HPV infection (2.68 versus 2.52; 2.62 versus 2.51; 2.50 versus 2.45; respectively, Wilcoxon-test, Figure 2B). We did not find other remarkable differences in *AICDA* gene expression or when we considered the AID-related mutations (Figure 2), suggesting that *AICDA* is rarely constitutively expressed and is rather transient at pan-cancer level. Interestingly, we found a positive correlation between AID-related mutations and the presence of M1 macrophages, T CD4 and CD8 cell populations (Figure 2D). In addition, in the vast majority of cases, the presence of these immune cell populations were co-expressed in the same cancer types in the same direction (Figure 2D). Inversely, *AICDA* gene expression was not co-expressed with these three immune cell types but was positively correlated with B cell naive and different properties of B cell receptor (BCR), as well as a lymphocyte cell infiltration signature (Figure 2E).

**Figure 2.**
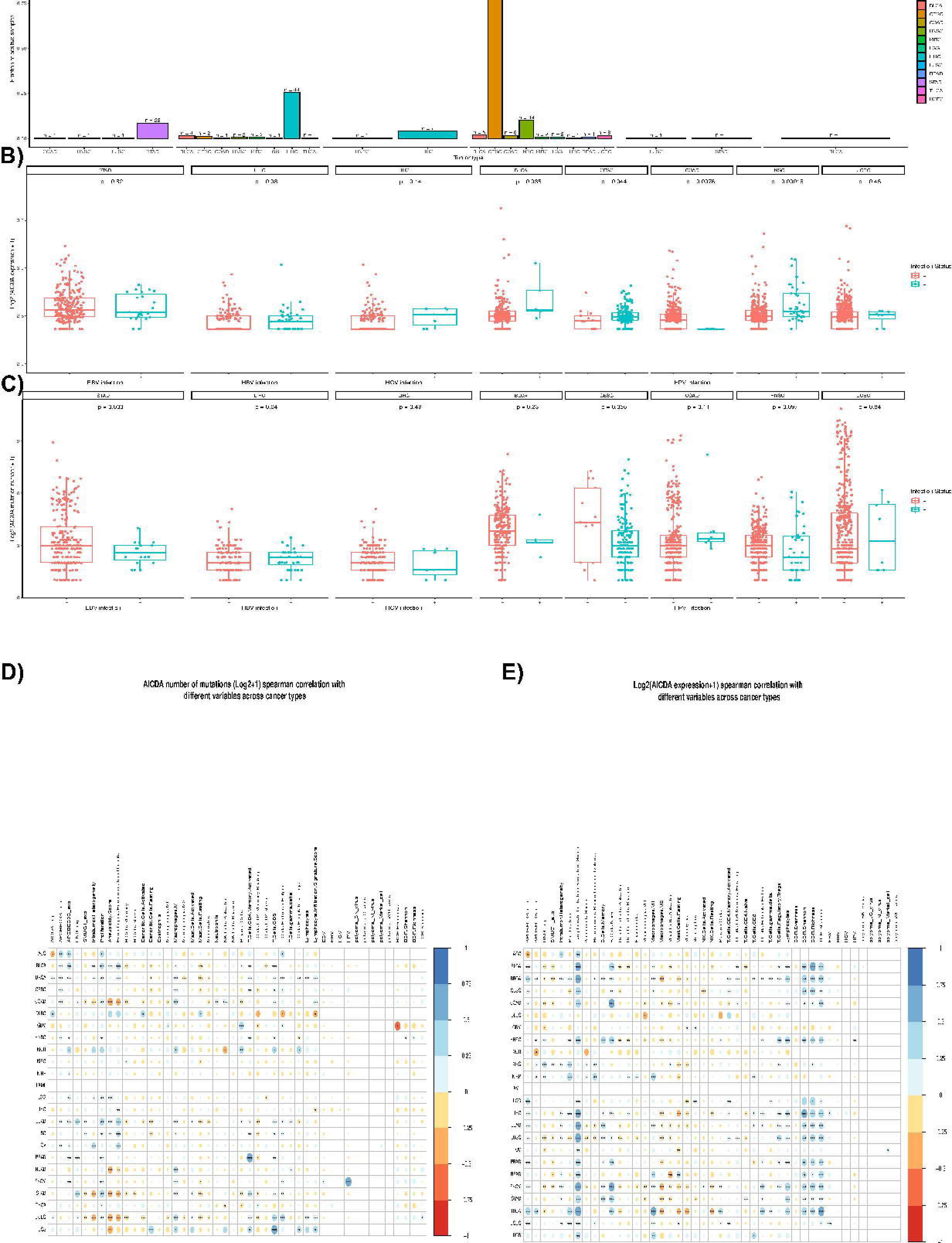
AID-mutations relation with immune features. Panel A, distribution of the presence of the genome of different viruses according to the cancer type. Panel B, distribution of the expression of *AICDA* according to the presence of different viral genomes per cancer type. Panel C, distribution of the number of AID-related mutations according to the presence of viral genomes per cancer type. Panel D, heatmap showing the correlation according to the number of AID-related mutations and the expression of different immune cell types, genes of interest or the presence of different viral genomes. Panel E, heatmap showing the correlation according to the expression of *AICDA* and the expression of different immune cell types, genes of interest or the presence of different viral genomes.

This suggests that only HPV infection, in general, does trigger *AICDA* expression but does not correlate with AID-related mutations.

### AID and APOBEC activity is higher at transcriptionally active domains but their relation with the MMR activity is contrariwise

Initial studies have revealed that the mutation frequency is increased in late-replicating regions (G2/M phases), mainly, due to increased MMR activity on early zones (G1/S phases; Tomkova et al., 2018). However, recent works have found 3D chromatin organization to be better correlated with mutational load than replication time alone but the direction, towards active or inactive domains, is shaped by the mutational signature (Akdemir et al., 2020). By using replication timing alone we found that the AID and SBS2 signatures have a clear late replication enrichment (global p-values = 3.09e^−35^ and 3.83e^−3^, respectively; Figure 3A). However, the enrichment zone changes across tumor types, being more "early" in Bladder-TCC, Cervix-SCC, Uterus-AdenoCa, Thy-AdenoCA, Lung cancers, and others which are mostly already reported APOBEC-prone cancer types (Supplementary Figure 11A-B). On the other hand, the SBS13 signature is not enriched globally (p-value = 0.88) but it is on early zone for the same tumors as SBS2, which is consistent with previous reports (Supplementary Figure 11C) (Morganella et al., 2016; Tomkova et al., 2018). In B-cells AID is mostly active in G0/G1 phase and has been proved to induce mutations before malignant transformation (Kasar et al., 2015). Alternatively, our findings indicate that it might change to be able to induce mutations at late replicating zones on hematological cancers and others (Kidney-RCC, Panc-Endocrine and Stomach adenocarcinoma; Supplementary Figure 11A). Next, we used topologically associated domains (TAD) boundary information of active and inactive domains, in terms of transcription, to see the distribution of AID/APOBEC mutations across chromating folding domains (Akdemir et al., 2020). We found AID mutations occuring more towards active domains than inactive (FC = 3.63; p-val = 5.01x10^-98^), specially at the TADs boundaries (Figure 3B). As previously described, we found that APOBEC signatures are also causing mutations towards active domains but the active/inactive ratio is notably higher for the SBS13 than the SB2, indicating distinct molecular underpinnings (Supplementary Figure 12).

**Figure 3.**
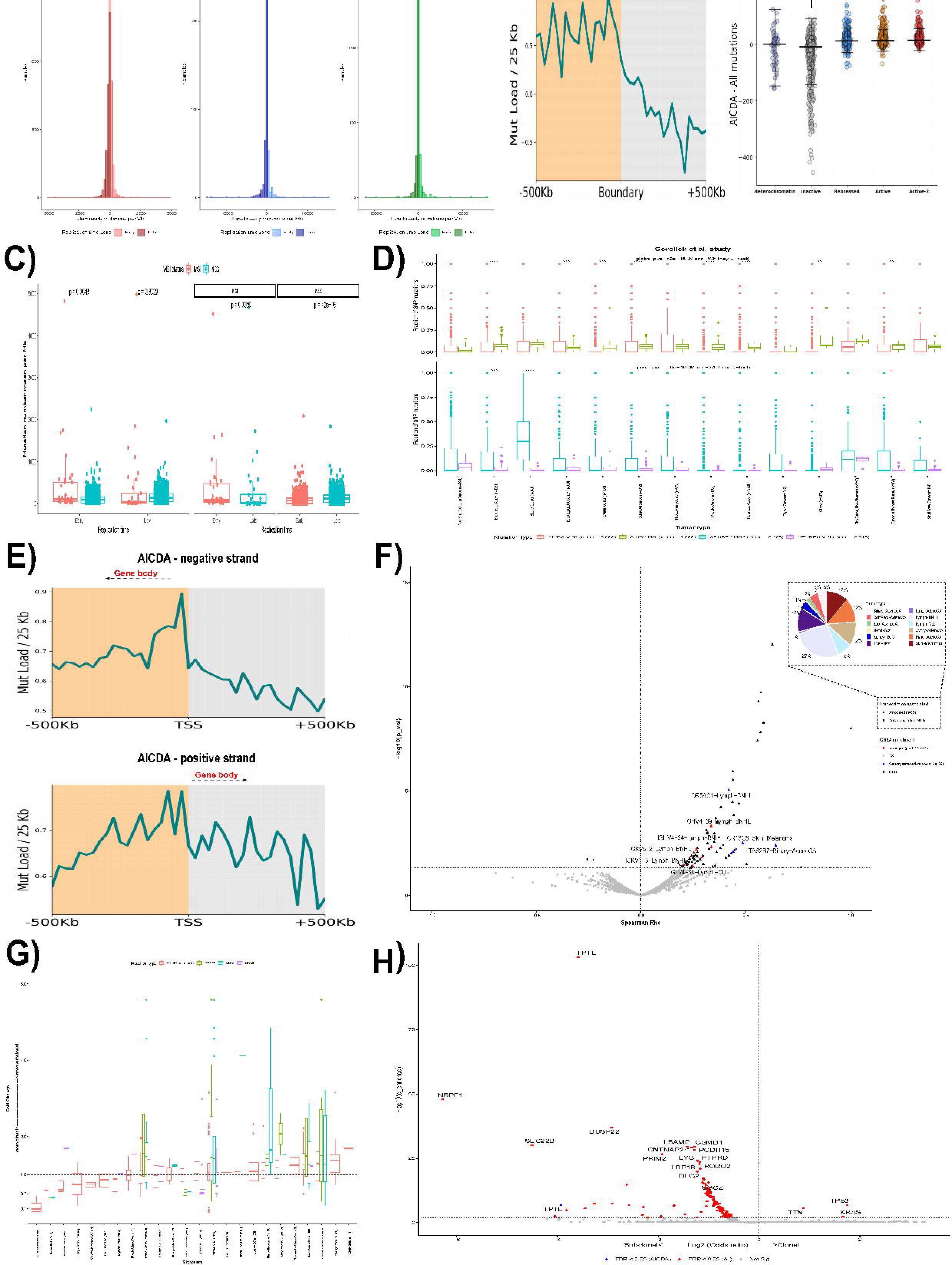
AID-mutations interplay with replication, transcription and clonality. Panel A, distribution of late to early replication time according to AID (left panel), and APOBEC signatures (SBS2 and SBS13), on the middle and right panels, respectively. Panel B, average profile of AID somatic mutations accumulation in 2,775 cancer samples and replication timing across 500 kb of TAD boundaries delineating active to inactive domains (left); dot plots representing the distribution of the mutations in different domain types (right; Wilcoxon rank-sum test). Panel C, distribution of the number of mutations associated with AICDA according to the replication time when considering MSS or MSI samples (panels at right). Panel D, distribution of mutations comparing MSS and MSI tumors according to AID mutations (top panel), showing different cancer types with higher frequency of AID mutations in MSI tumors. On the bottom, the distribution of mutations comparing MSS and MSI tumors according to APOBEC mutations, showing that only in endometrial cancer, bladder and cancer of unknown origin, there is a higher number of APOBEC mutations in MSS samples (MSKCC cohort, n = 19,936). Panel E, average profiles of AID induced mutations accumulation in 2,775 cancer samples across 500 kb of TSS for negative strand genes (top) or positive strand genes (bottom). Panel F, volcano plot (n = 1,130) showing the genes whose expression and mutations are correlated per tumor type (p-adj < 0.05, Spearman Rho > 0), where colors indicate genes enriched in a specific pathway by DAVID database analysis and the pie chart (inset plot) indicates the distribution across tumor types of the associated genes. Panel G, fold change of signature activities (subclonal to clonal) across tumor types for AID, APOBEC (SBS2 and SBS13) and SBS9 mutational signatures for samples with measurables changes in their mutation spectra (n = 404) where box plots indicate the first and third quartiles of the distribution, with the median shown in the centre and whiskers covering data within 1.5x the IQR from the box. Panel H, volcano plot showing the clonality enrichment per gene (FDR-adjusted from one-sided Fisher’s exact test) against odds ratio where color indicates enrichment for AID or global somatic mutations. All but panel D were produced using the ICGC cohort.

Since previous studies have demonstrated MMR activity producing 800 bp fragments prone to APOBEC mutagenesis meanwhile also being one of the repair mechanism of AID activity within B-cells, we attempted to see the differences of AICDA/APOBEC mutagenesis between MSI or MSS tumors (Mas-Ponte and Supek, 2020). When analyzing MSI tumors from ICGC, we observed higher AID induced mutations in early replication time (p-value = 0.0059; Wilcoxon test) which was also the case if compared to MSS tumors (p-value = 0.0043; Wilcoxon test; Figure 3C); but not in APOBEC related signatures (Supplementary Figure 12D). Furthermore, we also validated this hypothesis by analyzing 19,936 additional tumors (MSKCC cohort) finding a significant increment on the global number of AID or APOBEC mutations (p-value = 2e^-16^ & 6e^-10^; Wilcoxon test; Figure 3D) towards MSI or MSS tumors, respectively. Therefore, our analyses suggest that both APOBEC and AID mutations have higher preference at active domains and that the latter can be further repaired by the MMR machinery, rather than be enhanced by it.

### Oncogenic AID activity differs according to transcription direction

AID activity within the normal context takes place especially during transcription elongation, when the polymerase becomes stalled, and requires a licensing step to regulate over-activity which can be bypassed when abnormal high nuclear levels of AID are present. On the other hand, R-loops, a hybrid structure of B-form double-stranded DNA and A-form dsRNA, are formed during transcription which increases DNA exposure and has been linked to AID activity (Ginno et al., 2012; Methot et al., 2018). By using genomic coordinates of R-loop associated regions and the ICGC cohort (WGS data), we attempted to answer if within the tumoral context, AID mutations were more localized in or out these regions compared to either APOBEC mutations (COSMIC SBS2/SBS13) or other mutations. Surprisingly, only 0.18% (1130/629,871) of all AID mutations were “in R-loops” which was not significantly different to those caused by SBS2 (0.19%; 456/241,695; p-value = 0.37; two-sided Fisher exact test) or SBS13 (0.19%; 400/204,922; p-value = 0.15) (Supplementary Table 3). Overall, this suggests that within the oncogenic context, AID promiscuous activity is not related to R-loop formation.

Since the R-loop forming regions do not cover all the transcription start sites (TSS), we next analyzed the AID mutation’s distribution around the TSS as previous studies showed recruitment of AID to those sites (Methot et al., 2018). By dividing mutated genes based on strandness, we found a very particular pattern on the negative strand AID mutated genes compared to the positive strand. Mutations accumulate near the TSS and towards the gene body while maintaining a more constant mutational load compared to the opposite direction, the positive strand (Figure 3D) or even if compared to APOBEC signatures at either strands (Supplementary Figure 14A and B).

Next, we wondered if the AID mutations of a specific gene were produced during transcription of that same gene. To answer this we used 1,130 samples (comprising 24 tumor types) from which the mutations and expression data was available (ICGC cohort) and correlated the number of AID induced mutations occurring in gene i (AID_Muts_gi_) to the expression of the same gene i (Exp_gi_) within each tumor type, since the expression and mutations varies greatly in this context. Only 6.6% of the total different mutated genes (21,341) were expressed from which 6.0% (81/1,403, not repeated genes) were correlated with its corresponding gene expression per tumor type (p.adj < 0.05, Spearman Rho > 0); additionally, more than one third of these genes were found in hematological cancers (Lymph-BNHL and Lymph-CLL). Gene set enrichment analysis revealed immunoglobulin V region related genes (adjusted p-value = 1.2x10^-11^) within hematological cancers (Figure 3F). All together, our analysis suggests that AID activity is coupled to the transcription process with immunoglobulin genes in hematological cancers following the line of “normal” context as expression is more constitutive. However, the mutations in other genes are probably produced during short term transcription of both the affected gene and *AICDA* whose dynamics depend on the strand location of the gene and hence the direction of transcription.

### AID temporal mutations varies across tumor types and is independent of replication timing

Point mutations, within a cell, arising before a chromosomal locus duplication will give rise to different cell lineages compared to the mutations happening after the duplication. WGS data can be used to infer the number of allelic copies and hence the ratio of duplicated to non-duplicated mutations within a gained region can be used to estimate the time point when the gain happened and define variants as clonal (happening at early points) or as subclonal (happening at more recent time points) (PCAWG Evolution & Heterogeneity Working Group et al., 2020). By combining 13 million point mutations (SNVs only) and available copy number variations (CNV) data from the PCWAG dataset (2,707 samples), we evaluated the molecular timing of AID provoked mutations and compared to APOBEC signatures (SBS2 and SBS13) and the hypermutation attributed signature (SBS9). We found that 90.5% of mutations were clonal and 9.5% subclonal but when stratifying by APOBEC (SBS2 and SBS13) or AID signatures the proportion was slightly higher within subclonal mutations (2.8% versus 1.6% in SBS2, 2.5% versus 1.3% in SBS13 & 5.2% versus 4.4% in AID; P ∼0, respectively; fisher exact test; Supplemental Figure 14C). However, when looking only at samples with significant change within the mutational spectra of clonal versus subclonal mutations (404/2,707 samples; P < 0.05, Bonferroni-adjusted likelihood-ratio test), we found that even though AID signature is slightly more subclonal globally (median fold change = 1.06; IQR = 0.85-1.31), it has different temporal preference across tumor types. For example, there was a 9.6 fold change towards clonal mutations (IQR = 3.9-33.8) in Skin-Melanoma but a 1.3 fold change towards subclonality (IQR = 0.83-1.61) in Breast adenocarcinoma. On the other hand, we found more stable behaviours towards APOBEC (SBS2 and SBS13, more subclonal) or somatic hypermutation (SBS9, more clonal) associated COSMIC signatures across tumor types and in the same direction, as previously described (McGranahan et al., 2015; PCAWG Evolution & Heterogeneity Working Group et al., 2020) (Figure 3G). Furthermore, separating mutations by replication zones, globally and per tumor type, showed no evident change suggesting that this temporal preference is independent of the replication timing (Supplementary Figure 14D). Next, we checked if there were genes enriched for clonal or subclonal mutations globally or if AID provoked only, by calculating the odds ratio (OR) and corrected p-value of observed clonal/subclonal mutations versus the expected adjusting by various genetic covariates. When considering only OR, genes with AID induced mutations were more subclonal (OR < 1) than if considering all mutations (56.0% versus 41.9%; P = 2e^−16^, two sided two-sample Z-test).

However, only a limited number of genes emerged as significantly enriched (Figure 3H). Summing up, we found that for some cancers like skin melanoma AID-mutations probably contribute to oncogenesis at early times meanwhile for others it might promote oncogenic fitness by late mutations.

### AID synergizes initial hotspot mutations through late mutations on weakly functional alleles

Since recent studies have unraveled that composite mutations, pair of driver–driver, driver– passenger, or passenger–passenger mutations on the same gene, can synergize the functional impact compared to their single-mutated contrapart, we analysed the contribution of AID induced mutations within this phenomenon by analyzing 31,353 samples comprising 41 tumor types from the MSKCC (Figure 4). As previously described, using a panel of 353 oncogenes (168 genes) or tumor suppressor genes (TSGs, 185 genes), we found that composite mutations occur more frequently in TSGs than in oncogenes (12.2% versus 6.0% of all mutations; P = 2e^−278^, two sided two-sample Z-test) (Gorelick et al., 2020; Saito et al., 2020) but interestingly when separating by AID induced compared to those of other origin, we observed a global contribution to the composite mutations of 6.9%; furthermore, within oncogenes 9% consisted of at least one AID induced mutation, compared to 5% within TSGs (Figure 4B). We further verified that biallelic loss was also enriched for AID composite mutations, as it was reported from global composite mutations, within TSGs since there were more truncating variants compared to oncogenes (64% versus 8%; P ∼0; fisher exact test; Figure 4C). Next we calculated gene enrichment for AID composite mutations globally and per tumor type to discard that the observations were due to randomness by modeling the AID composite mutational burden as a function of genetic covariates (see Methods). Surprisingly, we found enrichment for six genes including *FGFR3* especially among HNSCC with 20% corresponding to AID composite mutations, and lower lineage-specific proportions for *EGFR* (8.9% in Glioma), *PIK3CA* (∼ 4% in Breast, Endometrial, Cervical and Skin cancers), *FBXW7* (∼ 7% in Colorectal and Esophagogastric cancers); *PTEN* (2.5 and 4% in Endometrial and Cervical cancers) but not *TP53* since it was present across different tumor types (Q < 0.01; Figure 4A and 4F, Supplementary Tables 4-5). We used a similar approach for residue’s enrichment to avoid missing residues not enriched at gene level, *PIK3CA E726* was the most enriched (q = 2.59e^-58^, Fisher’s exact test) followed by *TP53 R213*, *EGFR A289* and *PIK3CA R88* (Figure 4G, Supplementary Table 6). Since most found residues happened to be of lesser positive selection, we next checked the cumulative proportion moving from frequent hotspots (greatest positive selection) to less frequent ones finding that AID composite mutations are five times more likely to happen than AID singleton mutations (P = 2e-^109^, two-sample Z-test for equal proportion) which has higher than the fold change (FC) between composite mutants (other than AICDA) to singleton mutants (FC= 2.3; P ∼ 0). Furthermore, any AID mutation was absent from the highest positive selective hotspots (i.e. *KRAS G12, PIK3CA H1047, TP53 R273*) suggesting that AID mutations have preference towards weakly functional alleles after acquisition of high positive hotspots (Figure 4E, Supplementary Table 7). To further evaluate this hypothesis, we added the allelic configuration and clonality to subset to mutations arising from the same tumor cell population and retain molecular timing information, we observed that both globally (69% versus 31%, P = 7e^−4^, two-sided binomial test, Supplementary Figure 14A) and within AID composites (73% versus 27%, P = 0.03, two-sided binomial test, Figure 4D) the most frequent hotspot mutation occurs first and is followed by a synergizing second mutation but only within oncogenes, which was the case of the minor mutation *PIK3CA E726,* between the kinase and the PI3KA domains, that occurs significantly after (p = 0.039, one-sided binomial test) than other stronger mutations (i.e. *PIK3CA E542, PIK3CA E545* at helical domain or *PIK3CA H1047* at the kinase domain) (Figure 4I, Supplementary Table 8) and is a product of AICDA promiscuous activity. When looking only at phase-able mutations (without the molecular timing variable) we observed that 88% of composite mutations on *PIK3CA* occurs in cis from which 26% were AID provoked; other genes with high percentage of cis AID composite mutations were *EGFR, KMT2D* and *APC* (Figure 4H, Supplementary Table 9). Some *PIK3CA* composite mutations have already been proved to increase cell proliferation, tumor growth but also PI3K inhibitor sensitivity in human breast epithelial cell lines, but to the best of our knowledge it has not been linked to be product of AID activity (Saito et al., 2020; Vasan et al., 2019).

**Figure 4.**
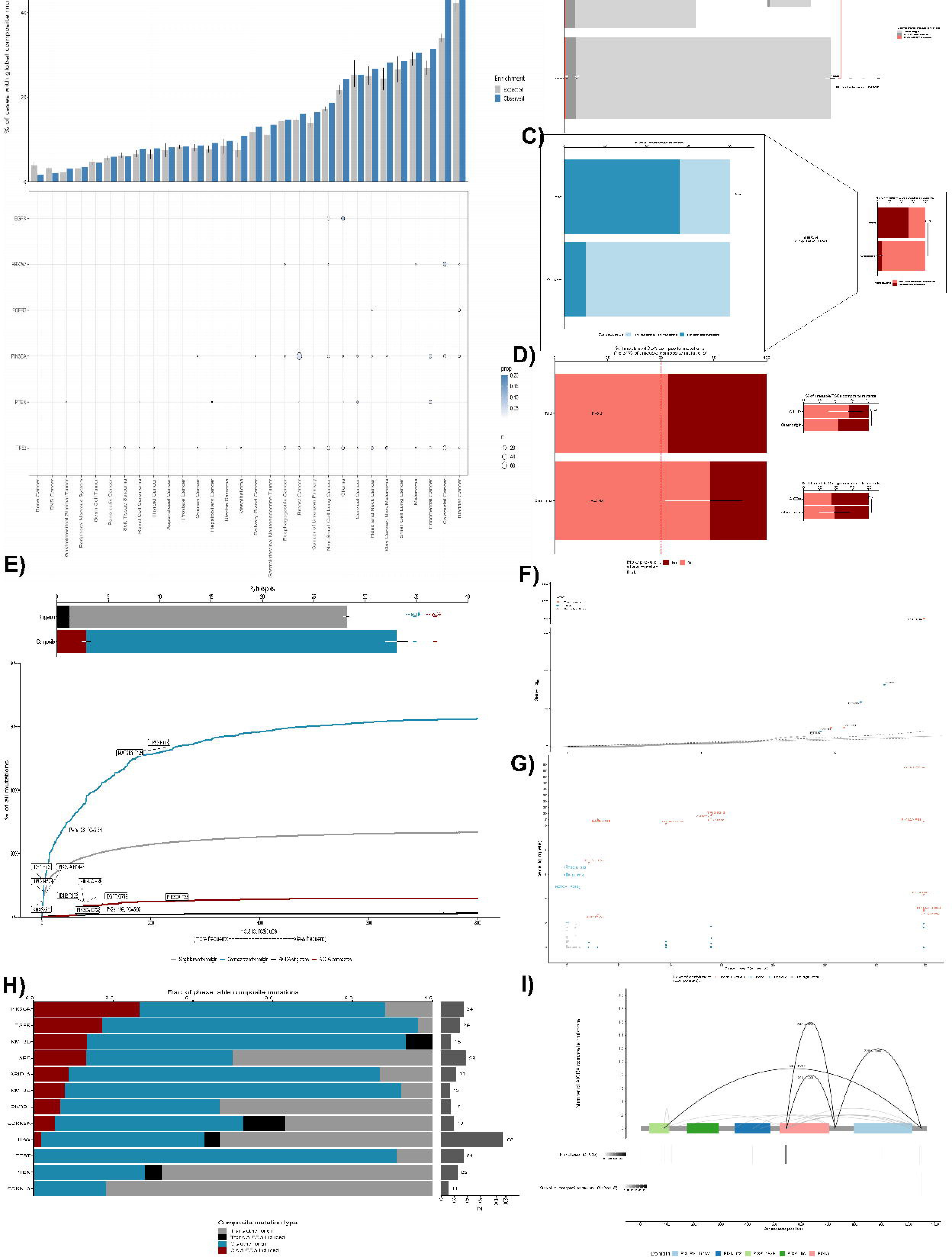
Impact of AID mutations on composite mutations. Panel A, AID composite mutations in enriched genes by lineage (n = 31,353 samples). Cases with global composite mutations and the expected value based on cohort size and mutational burden (top). Significant enrichment for AID composite mutations in cancer genes (FDR-adjusted P values from one-sided binomial test for enrichment) (bottom, n = 29,461). Panel B, percentage of total mutations (n = 130,962) that were composite by cancer gene function and composite mutation type (Two sided two-sample Ztest for equal proportions; color indicates the composite mutation type being compared; numbers on bars indicate absolute numbers of each mutation type). Subpanel B, relative contribution of at least one AID mutation within the composite or other origin to the composite mutations. Panel C, percentage of composite mutations by cancer gene function and consequence (Two sided Fisher’s exact test) in global composite mutations (n = 6,681, left panel) or only AID composite mutations (n = 472, right panel). Panel D, temporal order of acquisition of AID composite mutations by cancer gene function (n = 54 evaluable AID composites) using clonality and allelic configuration. Two-sided binomial test (left panel). Relative contribution of at least one AID mutation within the composite or other origin to each cancer gene function (Two sided Fisher’s exact test, right panel). Panel E, cumulative sum of the percentage of hotspot mutation utilization by decreasing frequency of population level hotspot mutations among composite or single mutations (AID or not AID provoked). Two-sided Mann–Whitney U test, fold-change (FC) of max composite to singleton values. Top inset, percentage of hotspots attributable to composite/singleton mutations (Two-sided two-sample Ztest for equal proportions, color indicates comparison for AID or not AID provoked). Panel F, significant enrichment for AID composite mutations in cancer genes (FDR-adjusted P values from one-sided binomial test for enrichment, n=29,461). Panel G, residue versus gene enrichment arising from AID composite mutations (FDR-adjusted from one-sided Fisher’s exact test for residues or one-sided binomial test for genes). Panel H, phase-able composite mutations by composite mutation type on different genes (left panel). Cases with phase-able composite mutations (right panel). Panel I, occurrence of PIK3CA AID composite mutations where arcing lines indicate the composite pairs (≥2 tumors, black color for AID enriched residues) and numbers indicate the amino acid position. Number of mutated cases at each individual residue and the significance value Q (FDR-adjusted P value from one-sided binomial test, bottom). Error bars in panels B-E indicate 95% binomial confidence intervals (CIs).

Additionally, we analyzed the contribution of other mutational processes to the composite mutations. Besides the aging signature, AID contributed more to the composite mutations than other signatures (Supplementary Figure 14B), opening the possibility of further research on the molecular implications of these mutations.

### The impact of AID-related mutations with ICI response

Because several recent studies pinpointed a potential role of APOBEC related mutations on the efficacy of ICI (Litchfield et al., 2021a; Wang et al., 2018), we sought to use the fraction of AID as a surrogate marker of ICI response. We used different available datasets analyzed (see Methods). We performed a random-effects meta-analysis comparing the overall survival (OS) of all these studies and comparing the impact of AID, to the APOBEC signature and the different single nucleotide variants (SNV). The details of this analysis are provided in the methods. Strikingly, the AID signature was associated with the best OS in all of the studies and the random-effects model showed also a favorable prognosis (median as the cut-off, Figure 5A). Moreover, the effect was still significant across almost all the studies independently of either decile chosen as cut-off at univariate (Supplementary Figure 15A) or multivariate adjusting for TMB (Figure 5B). Accordingly, the APOBEC signature was associated with a favorable prognosis, but not in all datasets. However, the random-effects model also indicated an overall favorable prognosis associated with APOBEC. The rest of SNV showed much more heterogeneous results and only T>A and T>G mutations were associated with favorable prognosis in the random-effects model (Figure 5A).

**Figure 5.**
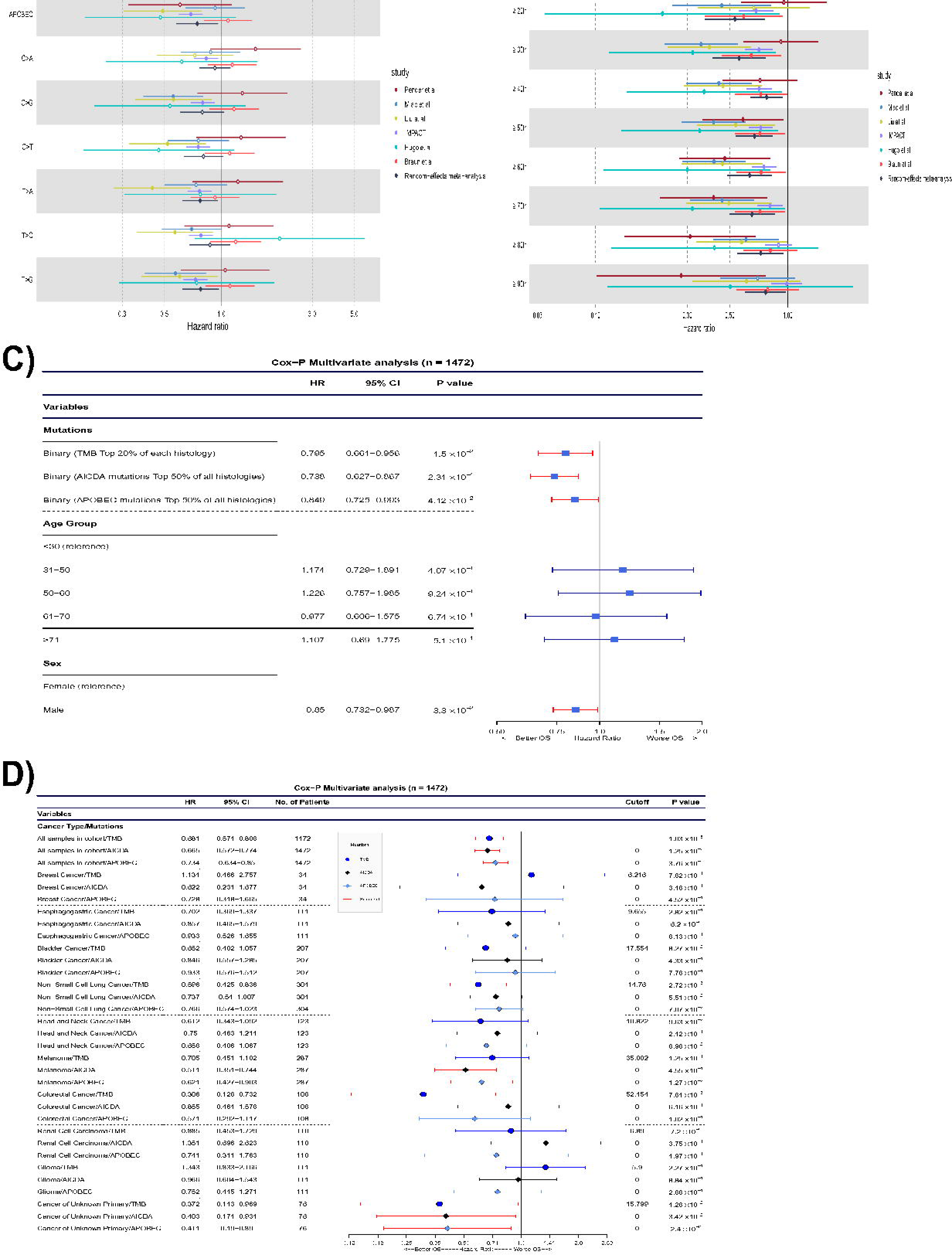
The impact of AID mutations on ICI response. Meta-analysis of the survival impact of the fraction of AID mutations in different studies. Panel A, effect of using AICDA/APOBEC (5th decile as cut-off) or SNV substitutions where AICDA remains significant across all the studies. Panel B, using all the deciles of the fractions of AICDA mutations (adjusting every decile of fractions of AICDA mutations per TMB ≥ 10 mut / Mb), the overall impact of AICDA with a better OS is present independently of the cut-off. Assessment of the prognostic value of the fraction of AICDA mutations in the IMPACT study. Panel C, forest plot of a Cox model of the global impact, after adjustment by TMB (top 20%), median APOBEC mutations, age and gender. Panel D, forest plot of the Cox model of the impact of AICDA mutations per cancer subtype.

Interestingly, within the largest study of IMPACT-MSKCC, the fraction of AID-related mutations (top 50% of all histologies as the cut-off) was also independently associated with both better OS (Hazard ratio [HR] = 0.715; 95% CI = 0.61−0.839; p = 3.81e^10−5^) and predictive value compared to TMB or APOBEC after adjusting by TMB (top 20% of each histology as the cut-off), APOBEC signature (top 50% of all histologies as the cut-off) age and sex (5C). It should be noted, that when using an univariate Cox proportional Hazards ratio model per every cancer type or adjusting by TMB ≥10, the results were also similar in the overall population of this study, but the clinical impact of AID-related fraction of mutations was only found in metastatic melanoma and cancer with unknown primary (Figure 5D; Supplementary Figures 15B and 15C). Additionally, there was practically no correlation between the fraction of AID mutations with the APOBEC signature neither globally nor by tumor type in this cohort and in the ICGC and TCGA datasets (Supplementary Figures 15D-F). Similarly by using four additional studies across different tumor types, we also found an association of high AID mutations with improved OS after adjusting by age, gender and TMB using the multivariate Cox model (Hugo et al., 2016a; Liu et al., 2019; Miao et al., 2018b; Pender et al., 2021b).

Overall, all the studies confirmed the independent prognostic value of the high fraction of AID mutations according to the median in the univariate and multivariate analyses.

### Landscape of AID-related neoepitopes and its relation with ICI response

Having found an association between AID activity and ICI benefit, we hypothesized that AID mutations might generate highly immunogenic neoepitopes. We addressed this by analyzing the neoepitopes that were products of AID activity on the TCGA cohort and on melanoma patients treated with Nivolumab (anti-PD-1) (Riaz et al., 2017). A recent bioinformatic-experimental study using immunogenic and non-immunogenic peptides, experimental testing and X-ray structures showed that TCR binding and recognition improves with the presence of hydrophobic amino acids (aromatic W, F, Y followed by V, L and I) at specific “MIA’’ positions (position P_4_-P_1-Ω_) due to increased structural avidity, stacking interactions, hydrogen bond acceptance and limited rotational freedom with the TCR (Schmidt et al., 2021). Additionally, as a previous study showed that APOBEC promiscuous activity increases neopeptide hydrophobicity (Boichard et al., 2018), we wondered if AID-related mutations led to the production of not only more hydrophobic neoepitope but more “Immunogenic” in terms of amino acid changes (W, F, Y, V, L, I over others) at MIA positions and if these effects were different due to clonality, histology or mutational processes. We computed the PRIME %rank score and used it to classify neoepitope as “Immunogenic” or “Non-Immunogenic” (see Methods), on a list comprising 2,143 patients (TCGA) from which RNA-seq, HLA haplotyping, clonality and mutational process origin data was correctly assessed; we also restricted the analysis to only patients with >1 FPKM expression on the genes originating the neopeptide, microsatellite stability and intact antigen presentation related genes. We analyzed 286,909 neoepitope from which only 17.75% were predicted to be immunogenic but interestingly they occur more frequently within clonal neoepitope than in subclonal (38% versus 30%, P = 1x10-^27^; two sided Fisher-exact test; Figure 6A). When accounting mutational signatures and dividing by histology, we observed a global trend towards higher number of immunogenic clonal neoepitope (ICNs), compared to subclonals, which reached significance for signatures characteristic of a cancer type, i.e. smoking-associated signatures in LUAD and LUSC or mismatch repair (MMS) signatures in HNSC, LUSC and BLCA; additionally, AID ICNs were significantly higher in BLCA and LUSC, meanwhile APOBEC in BLCA and CESC (Supplementary Figure 21A). Moreover, when comparing the number of ICNs produced by different mutational processes within each tumor type, we observed that in general, there was a higher number due to other signatures different to AID, especially APOBEC or smoke-associated signatures (Supplementary Figure 21B). Because our results suggested higher presence of immunogenic neoepitope, in terms of numbers, provoked by mutations occurring earlier which is unrestrained of mutational origin, we restricted the subsequent analyses to only ICNs.

**Figure 6.**
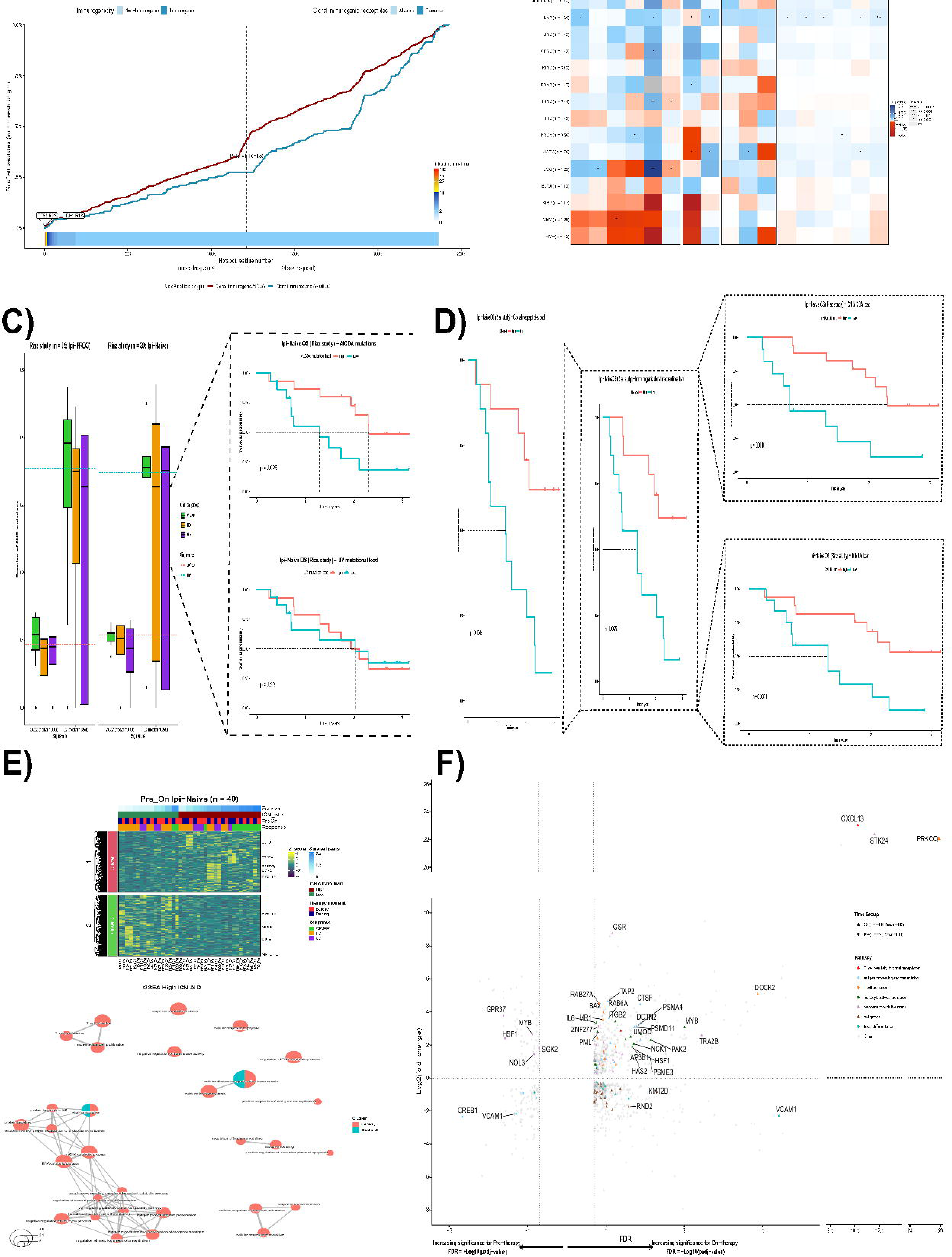
Landscape of AID-related neoepitopes and its relation with ICI response. Panel A, percentage of neoepitopes originating from clonal/subclonal mutations in which color indicates comparison for immunogenic or non-immunogenic (top left; Two-sided two-sample Ztest for equal proportions), calculated by Prime. Top right shows the comparison of the percentage of samples having at least one AICDA ICN versus APOBEC ICN (“Presence”; Two- sided two-sample Ztest for equal proportions). Bottom plot shows the cumulative sum of hotspot mutation utilization that gives rise to ICN by decreasing frequency of population level hotspot mutations due to AICDA or APOBEC signatures (Two-sided Mann–Whitney U test, FC of median AICDA to APOBEC values). Panel B, heatmap of gene expression comparison between ICN AICDA “Presence” versus “Absence” groups across tumor types/all tumors (n = 2,143; two-sided Wilcoxon test) measured as log2 FC. Panel C, AICDA and UV related mutations are higher within responders (CR/PR) compared to non-responder (SD or PD) in both Ipi-Prog and Ipi-Naive cohorts (left, data are presented as median and interquartile range) but OS only associates with AICDA mutations within Ipi-Naive patients (right). Panel D, OS prediction within Ipi-Naive patients improves when using ICN AICDA load (top right), ICN UV load (bottom right), ICN load (middle) or clonal neoepitopes load (left), lowest to highest log-rank p-values. Panel E, heatmap and hierarchical clustering of DEGs between pre and on-therapy grouped according to high/low ICN AICDA load (top). GO enrichment analysis (q < 0.05) of the gene clusters found in top E. Panel F, DEGs (p-adj <0.20) between high ICN load patients versus low ICN load for pre-therapy, where increasing negative values on the x axis shows higher significance (+Log10[p-adj]), or on-therapy, where increasing positive values on the x axis means higher significance (-Log10[p-adj]). The Y axis shows upregulated (FC > 0) or downregulated genes (FC < 0) and colors indicate genes enriched in a specific pathway by GO analysis. Panels A and B correspond to TCGA cohort (n = 2,143) meanwhile panels C-F to ICI treated melanoma cohort (Riaz et al., n = 68).

Strikingly, albeit a global higher number of APOBEC induced ICNs was present, the proportion of samples having at least one ICN produced by AID (classified as “Present”) was practically three times higher than those provoked by APOBEC globally (32% versus 11%, P = 1x10^-65^; two sided Fisher-exact test; Figure 6A). We next sought to compare the cumulative distribution of the AID/APOBEC ICNs in terms of population hotspot mutations recurrency finding that AID produces ICNs at hotspots with greater positive selection (FC = 1.59; P = 3e-^41^, two-sample Z-test for equal proportion, Figure 6A) which could give rise to higher possibilities of immune recognition and improved tumor control. By comparing tumors harboring at least one AID ICN (“Presence”) to those which did not (“Absence”), we found an increased fraction of CD8, CD4 memory activated and follicular helper T-cells that were “exhausted” by higher expression of the inhibitory immune checkpoint molecules PD-1, PD-L1, PD-L2, CTLA-4 and LAG3. Furthermore, these observations were seen in the majority of tumor types but the increment was only significant when accounting for all the samples (n = 2,143) or for LUAD (Figure 6B, two-sided Wilcoxon test).

Since these findings suggested that AID mutations inducing ICN as possible explanation of ICI response, we next analyzed a cohort of 68 melanoma patients treated with anti-PD-1 (Nivolumab) from which WES, neoepitopes and RNA-seq data was available prior treatment (pre) or 4 weeks after initiation of Nivo (on) (Riaz et al., 2017). Through all the analysis we separated patients as Ipi-Prog (n = 35), which had previously progressed on anti-CTLA-4 treatment (Ipilimumab), or as Ipi-Naive, which only received Nivo (n = 33). First, we looked at the distribution and effect of AID mutational load on survival compared to UV related mutations. We found that the responders (CP/PR) had a higher number of AID-related mutations compared to the PD or SD groups (Ipi-Prog median = 0.094; Ipi-Naive median = 0.108). However, it was not significantly different and was also observed for UV mutations (Ipi-Prog median = 0.354; Ipi-Naive median = 0.349). Conversely, the effect on OS was markedly different being associated with prognosis only when using AID mutations within Ipi-Naive patients (log-rank p = 0.026) but not with UV mutations in neither naive or progressive patients (log-rank p = 0.93 & p = 0.34; Figure 6C and Supplementary Figure 22A). The AICDA ICN load improved survival prediction better (log-rank p = 0.0016) than if using global clonal neo-epitopes load (log-rank p = 0.0042), global ICN load (log-rank p = 0.0025) or UV ICN load (log-rank p = 0.0071; Figure 6D). As the effect was tightly marked only in Ipi-Naive patients, we focused the subsequent analysis on only this group.

When coupling RNA-seq data (n = 20), we found 64 upregulated and 110 downregulated genes comparing patients with high ICN AID load versus low within pre-therapy samples (q < 0.20; Supplementary Table 10). Gene Ontology (GO) analysis identified down regulation of antigen presentation and TNF signaling pathways (q-value <0.05; Figure 6F & Supplementary Figure 22B). We also observed an increased expression of the inhibitory immune checkpoint molecules PD-1, PD-L1, PD-L2, CTLA-4, ICOS, LAG3 and cytolytic activity (Supplementary Figure 22C and D). These results are consistent with both our previous analysis on TCGA data and previous studies (McGranahan et al., 2016; Riaz et al., 2017; Van Allen et al., 2015).

Next we endeavored to identify expression changes on patients that responded (according to ICN AICDA load) after 4 weeks of Nivo treatment by comparing pre-therapy to on-therapy data from the patients (n_pre_ = 20; n_on_ = 20). From the 811 genes, found to be differentially expressed (q < 0.20; Supplementary Table 11), 404 were up-regulated and involved in antigen processing and presentation, T-cell activation (e.g. *PRKCQ, CD8B, CD38, CD151, MALT1*), leukocyte cell-cell adhesion, response to oxidative stress (*STK24*, *GSS, GCLC, PDK1*) and T-cell reactivity to clonal neoepitopes (CXCL13 and CCR5) (q-value <0.05), the last ones being recently described (Litchfield et al., 2021b). On the other hand, downregulated pathways included mainly (407 genes; q-value <0.05) cell growth, B-cell differentiation and some chemokines (*CXCL11, CCL4* and *CCL14*) or chemokine receptors (*CCR3* and *CCR8*) (Figures 6D and 6F). Furthermore, we also observed an increased expression *CXCL13* of and *CCR5* on the TCGA samples with high ICN AID load (Figure 6B). Altogether, these results show possible explanations of why AID mutations reflect a more straightforward approach to predict response to ICI naive treatment.

### scRNA analysis reveals activation of *AICDA* expression under oncogenic conditions

To better understand *AICDA* expression across tumor types we looked at their normal counterparts by analyzing scRNA-seq data from ∼ 600, 68 and 350 thousand cells from the human cell landscape (HCL), peripheral blood mononuclear cells (PBMCs) and the mouse cell atlas (MCA), respectively (Han et al., 2018a, 2020a; Zheng et al., 2017b). When analyzed by tissue, we found that *AICDA* expression occurs mainly in adult epityphlon and adult pleura (Figure 7A) in human samples and in adult small intestine, ovary, pleura and spleen in mouse samples (Figure 7B). The expression was significantly stronger in adult stages than in fetal or embryonic stages (Figure 7D). Additionally, when using the cell type annotation, in the HCL, we observed the highest expression in B-cells followed by fibroblasts (Figure 7A). Further analysis of only PBMCs led to the finding that *AICDA* is also expressed in CD8 T-cells and induced regulatory T-cells, but at lower levels than that of B-cells (Figure 7C). Interestingly, the expression of the base excision repair (BER) gene *UNG* and the MMR genes *MSH2/MSH6*, involved in downstream reparation of AID mutations, were expressed altogether with *AICDA* only within B-cells, CD8 T-cells and induced regulatory T-cells (Tregs).

**Figure 7.**
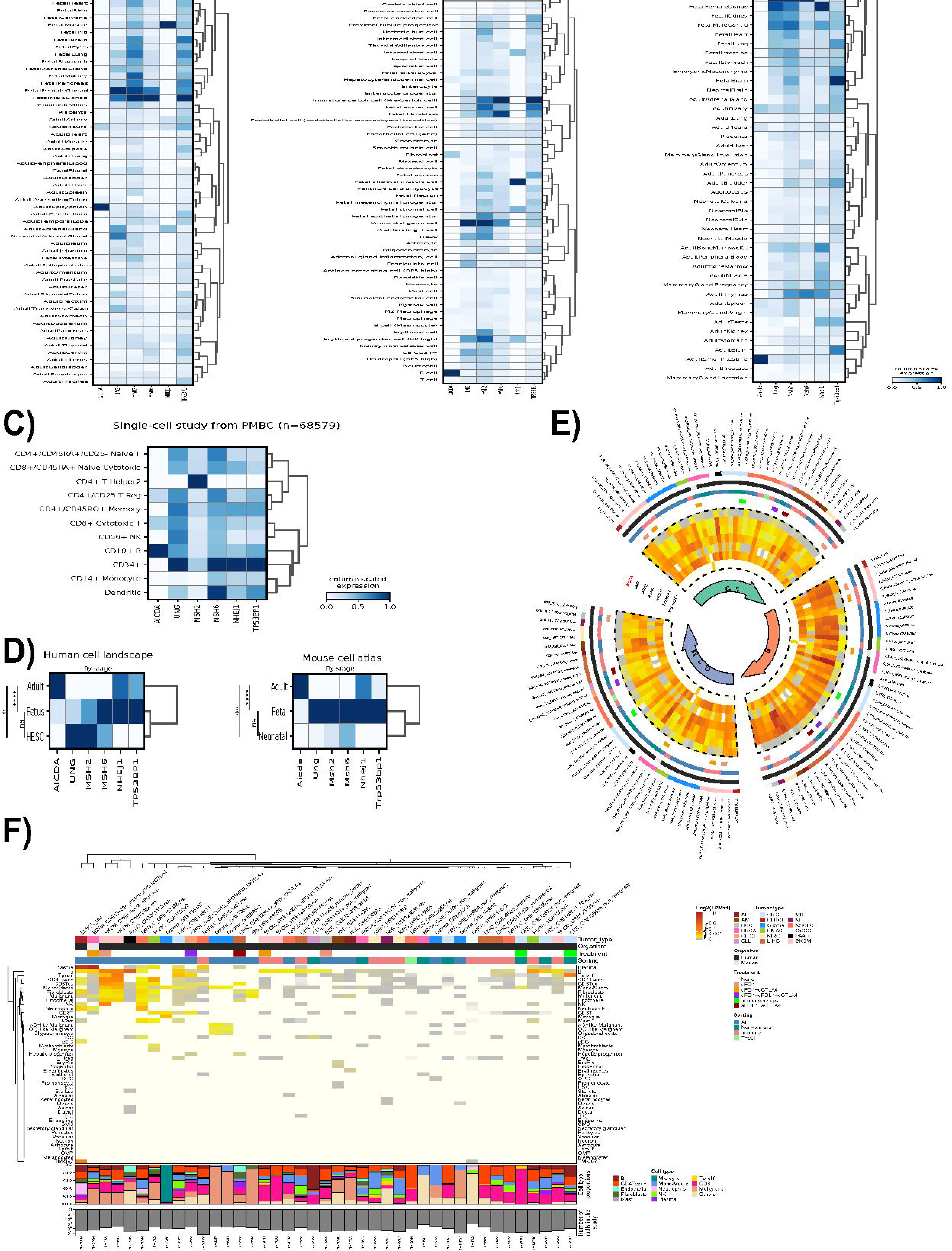
scRNA analysis reveals activation of *AICDA* expression under oncogenic conditions. Expression heatmaps of *AICDA* and genes involved in the repair of *AICDA*-related mutations. Panel A, human cell landscape (HCL) showing *AICDA* expression mainly in adult epityphlon and adult pleura tissues (left) or in B-cells and fibroblasts when accounting for cell subtypes (right). Panel B, mouse cell atlas (MCA) showing *AICDA* expression mainly in the adult small intestine, ovary, pleura and spleen. Panel C, normal PBMCs showing *AICDA* expression only in B-cells, CD8 T-cells and induced regulatory T-cells. Panel D, *AICDA* is significantly more expressed (Wilcoxon two-sided test) in the adult stage than in fetal or embryonic stages in both human (left) and mouse (right). Panel E, expression heatmaps of *AICDA* and genes involved in the repair of *AICDA*-related mutations as a function of the cell cycle stage across different oncogenic single cell studies (n = 40). *AICDA* expression is slightly higher in the G2M phase and the repair genes are more expressed during S phase; additionally, expression is dropped when tumoral cells are depleted by sorting. Panel F, *AICDA* expression heatmap as a function of the cell subtypes across different oncogenic single cell studies (n = 41; top), barplot of cell subtypes proportions (middle) and number of cells in each study (bottom). *AICDA* is expressed in malignant cells in most tumor types but only in some for the immune and stromal cell populations.

Having in mind deciphering the cell subtypes’ contribution to *AICDA* expression and its regulation through the cell cycle, we gathered information from 41 oncogenic single cell studies comprising around 1.5 million cells and 18 tumor types. Malignant cells express *AICDA* across all but one tumor type (BCC), being stronger within SKCM, MB, NSCLC, HNSC and Glioma. Moreover, the expression was lost when depleting the samples of tumoral cells and was not present in Tregs. Interestingly, a subset of the immune population cells, especially B-cells and exhausted CD8 T-cells within SKCM, DLBCL and NSCLC, and monocytes/macrophages in some SKCM and some Glioma studies; along with fibroblasts and endothelial cells from the stromal population, also express *AICDA* (Figure 7F). Surprisingly, its expression was observed across all cell cycle phases. However, it was slightly higher at G2/M; however, BER and MMR related genes are markedly more expressed at S phase, meanwhile the expression of nonhomologous end joining (NHEJ) related genes (*NHEJ1, TP53BP1 or Trp53bp1* in mouse) remained practically unchanged. Additionally, the expression was higher in SKCM, DLBCL and Gliomas independently of treatment (Figure 7G) . Altogether, these results suggest that cells activate *AICDA* expression after malignant transformation, which is independent of the cell cycle phase. However, the level of *AICDA* expression varies upon tumor type.

## Discussion

By integrating more than 50.000 bulk level samples and 2.5 million cells at single cell resolution across 80 tumor types and different data levels, we present, to the best of our knowledge, the first study shedding light into the oncogenic and clinical implications of AID at pan-cancer scale. Our results point to the idea that *AICDA* expression, which is activated after malignant transformation, is no longer tied to the cell cycle regulation and, albeit transient, it induces traceable mutations with important functional and clinical implications that are mainly produced during the transcriptional activity of the mutated gene. Firstly, our single-cell RNA-seq analysis revealed that only a selected number of tissues (e.g. adult pleura, small intestine) express *AICDA* under normal conditions which is consistent with previous studies at bulk level (Lonsdale et al., 2013; Uhlen et al., 2015); however, after acquiring malignancy the expression is seen independently of tissue origin as already reported for some tumor types (Kasar et al., 2015; Komori et al., 2008; Li et al., 2019b; Lossos et al., 2004; Nonaka et al., 2016; Sawai et al., 2015; Shimizu et al., 2014). Moreover, our single cell data analysis shows that B-cells within the TME of some cancer types are expressing *AICDA* which correlates with our findings on the TCGA data showing correlations of *AICDA* expression with the B-cell and lymphocyte infiltrate populations.

It was previously reported that in germinal center B-cells the *AICDA* expression is higher at G2/M cell cycle phases but its mutational effects are mostly active in early G1 (after being transported to the nucleus). However, this regulation is lost in lymphoma cells (Milpied et al., 2018; Wang et al., 2017). On the other hand, *UNG* expression is most abundant at S phase meanwhile *MSH2/MSH6* at G1/S, both implicated in AID induced mutations’ repairment (Álvarez-Prado et al., 2018; Delgado et al., 2020; Kasar et al., 2015). The oncogenic single-cell datasets analyzed proved that this cell cycle regulation is globally lost in all tumor types and that cells at G2/M phases are more susceptible to AID promiscuous activity given that: i) *AICDA* expression is higher; ii) the BER and MRR related genes are less expressed and iii) global transcriptional activity is also increased. This hypothesis is supported by our findings, using WGS data, that AID mutational load is increased: at transcriptionally active TAD domains (compared to the background), close to TSS and in MSI tumors. Regarding the different AID mutational behaviour depending on the strand location of the gene, we propose a model were the negative strand is more prone, than the positive strand, to AID attack at naked transcribed breathing dsDNA (normally located near TSS) and is followed by attack at DNA stem-loops and transcription bubbles (but not at R-loops) being generated as the RNA polymerase transcribes (Branton et al., 2020).

Summing up to those findings plus that *AICDA* expression and related mutations were not correlated in the TCGA nor in ICGC datasets and that only expression but not the mutations correlated with viral infection in some cancers, it is tempting to speculate that the genotoxic effect of AID might be due to short term activation of *AICDA*, which have been seen in APOBEC (Langenbucher et al., 2021). Indeed, in a fate mapping study, *AICDA* expression was present in a fraction of non-lymphoid embryonic cells (Rommel et al., 2013). Furthermore, *AICDA* transcripts in lymphocytes have a half-life of only one hour (Dorsett et al., 2008), supporting the lack of correlation between AID-related mutations and *AICDA* expression.

Despite, ephemeral, *AICDA* expression mutational footprints are widespread across cancers, and presumptively across mammals, with similar mutational frequency compared to APOBEC but higher contribution to driver oncogenes, to composite mutations and to the production of higher quality neo-epitopes. Already reported AID off-target activity, outside lymphomas, is limited especially to *TP53, KRAS and MYC* in gastric, colorectal and skin melanoma (Li et al., 2019b; Nonaka et al., 2016; Shimizu et al., 2014). We thoroughly extended this data and found that AID activity have preference towards least positive selection hotspots that synergizes with previous stronger hotspot mutations; this is the case for the minor mutation PIK3CA E726, especially present in SKCM and BRCA, that might confer higher PI3K inhibitor sensitivity (Saito et al., 2020; Vasan et al., 2019).

Finally, we found that AID-related fraction of mutations is an independent prognostic value to ICI response using > 2,000 samples even after adjusting by TMB. AID-related neoepitopes exhibited distribution towards clonal hotspots with greater positive selection which could result in improved immune recognition; however, this is avoided by tumor induced immune exhaustion. It should be noted that the statistical power in individual histologies is reduced, and as sample sizes increase, additional histology-specific associations may appear in future larger prospective studies that may lead to a formal validation of the predictive value of AID-related signature on ICI response and the results regarding the AID-related neoepitopes. It is also important to highlight that there could be some analytical bias related to the combination of different datasets using different mutation calling approaches. However, the signal associated with the AID-related mutations was similar throughout the studies and the pipelines and results of the different included studies are public and well standardized, limiting in part this mutation call bias.

We propose a model in which AID ICN have higher probabilities of being recognized by T-cells, triggering selective expression CXCL13, previously found to be a marker of antigen reactive CD8 T-cells, for recruitment of CXCR5+ T and B cells (Litchfield et al., 2021b). These recruited cells, subsequently exhausted by continuous expression of inhibitory immune checkpoint molecules, can be reinvigorated after ICI treatment.

Overall, we pieced together an immense part of the oncogenic AID puzzle but many parts still need to be found, especially filling gaps with biological validations as the results, here presented, holds the promise of important clinical applications.

## Acknowledgments

We greatly thank all investigators, funders, and industry partners that supported the generation of the data within this study, as well as patients for their participation. Specifically, we thank Eli Van Allen, Roel Verhaak and Timothy A Chan for the academic datasets.

The results published here are in whole or part based upon data generated by the TCGA Research Network: https://www.cancer.gov/tcga and the ICGC consortium: https://dcc.icgc.org. Graphical abstract created with biorender.com.

## Author contributions

I.H.-V. and A.A. designed the study. I.H.-V., K.M., K.H.-X.,M.P., F.B., M.T., A.I., A.D.,M.S. and A.A. performed clinical work. I.H.-V., K.C.A., D.R., G.C., K.L. and A.A. analyzed data. I.H.-V., K.C.A.,D.R.,G.C., A.D., M.S. and A.A. interpreted data. I.H.-V. and A.A. wrote the manuscript.

## STAR METHODS

## KEY RESOURCES TABLE

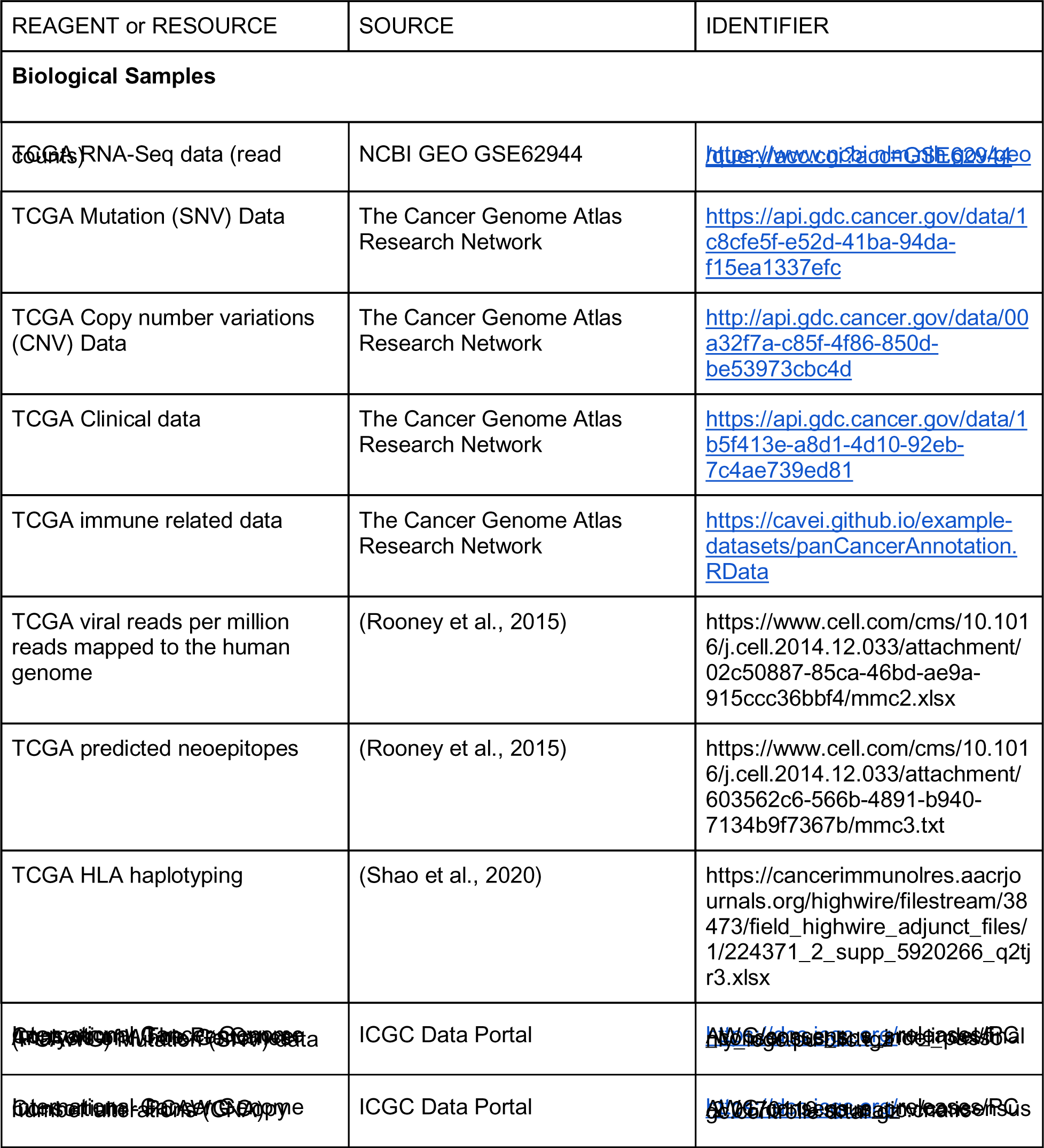
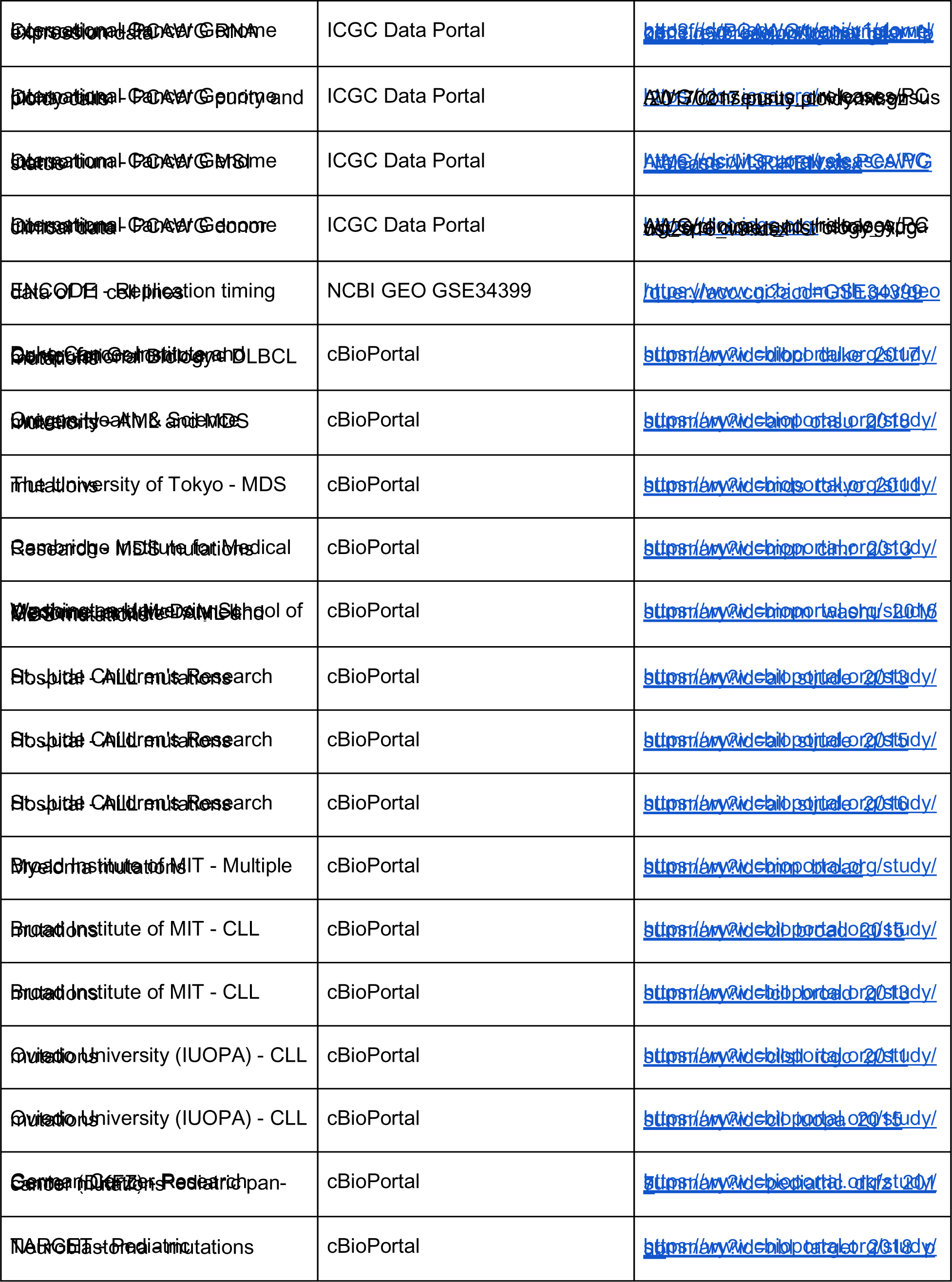
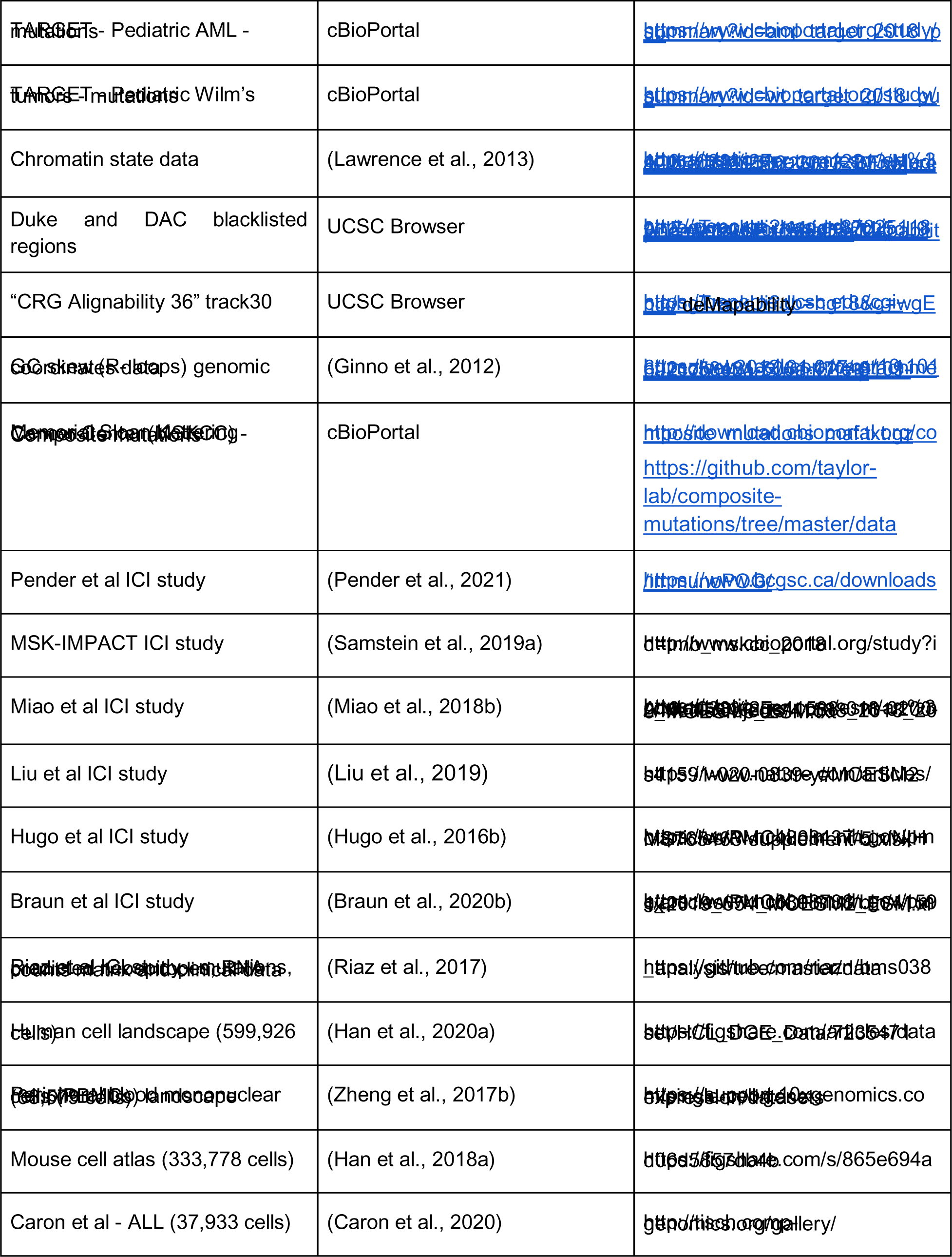
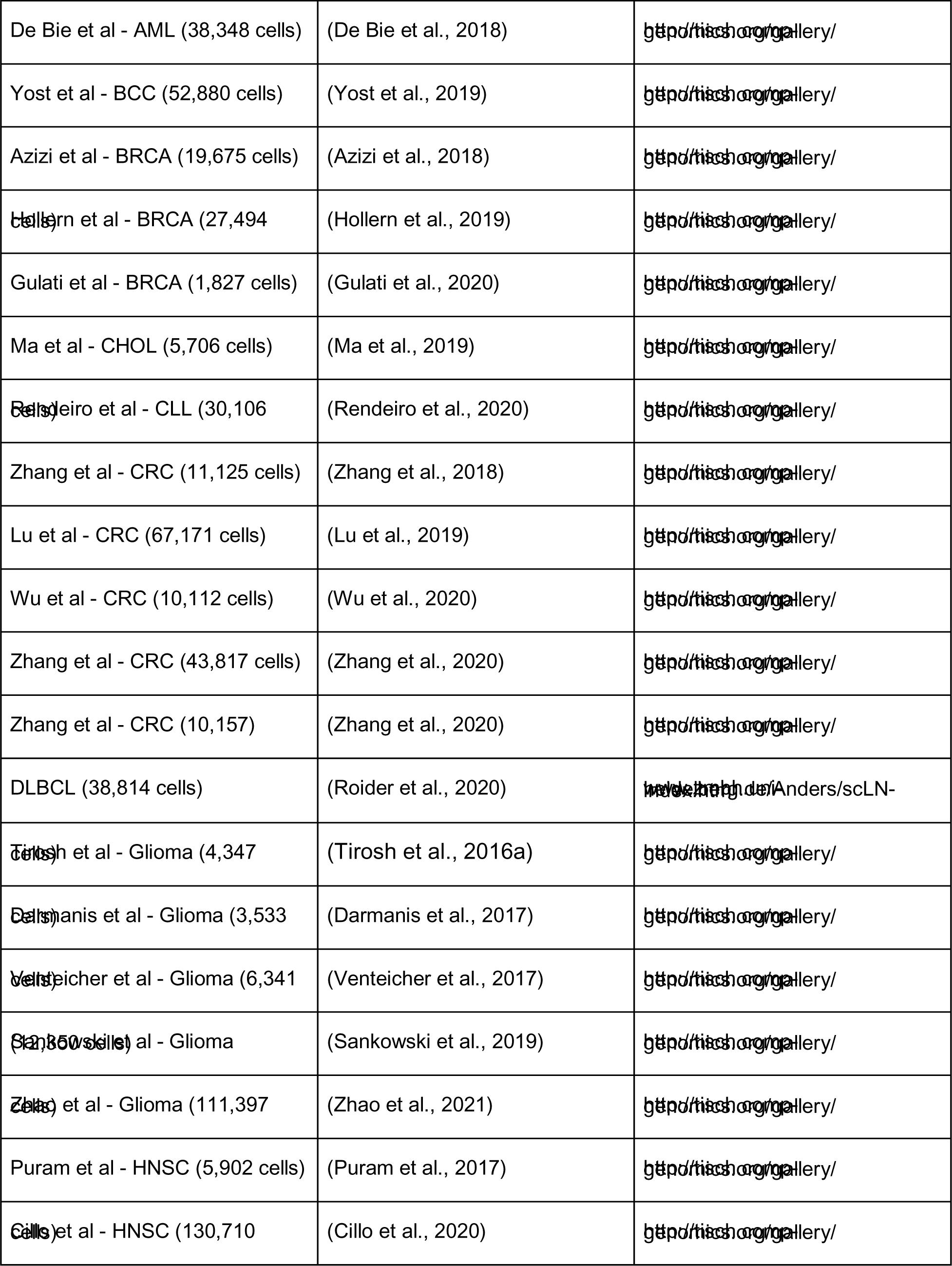
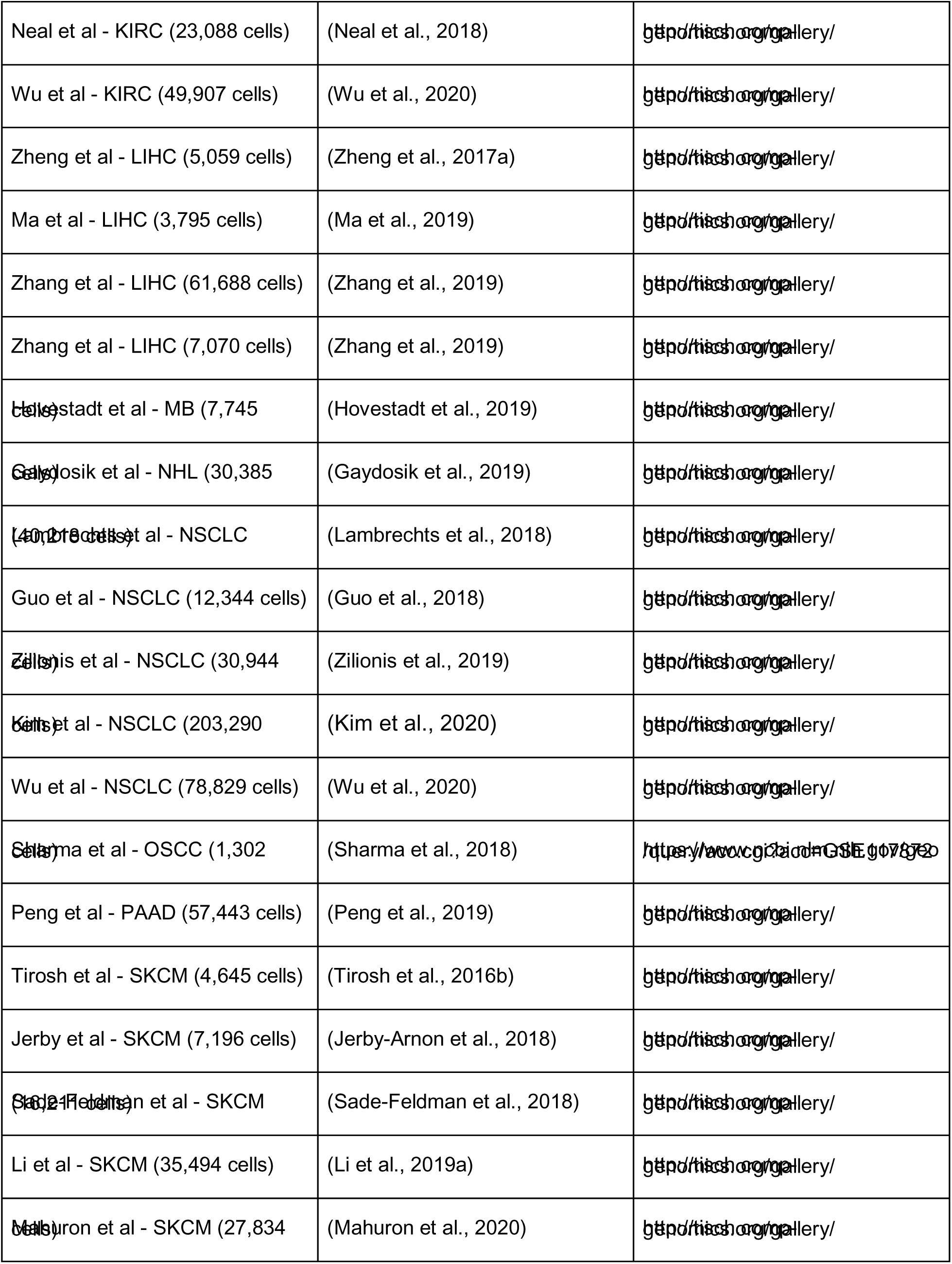
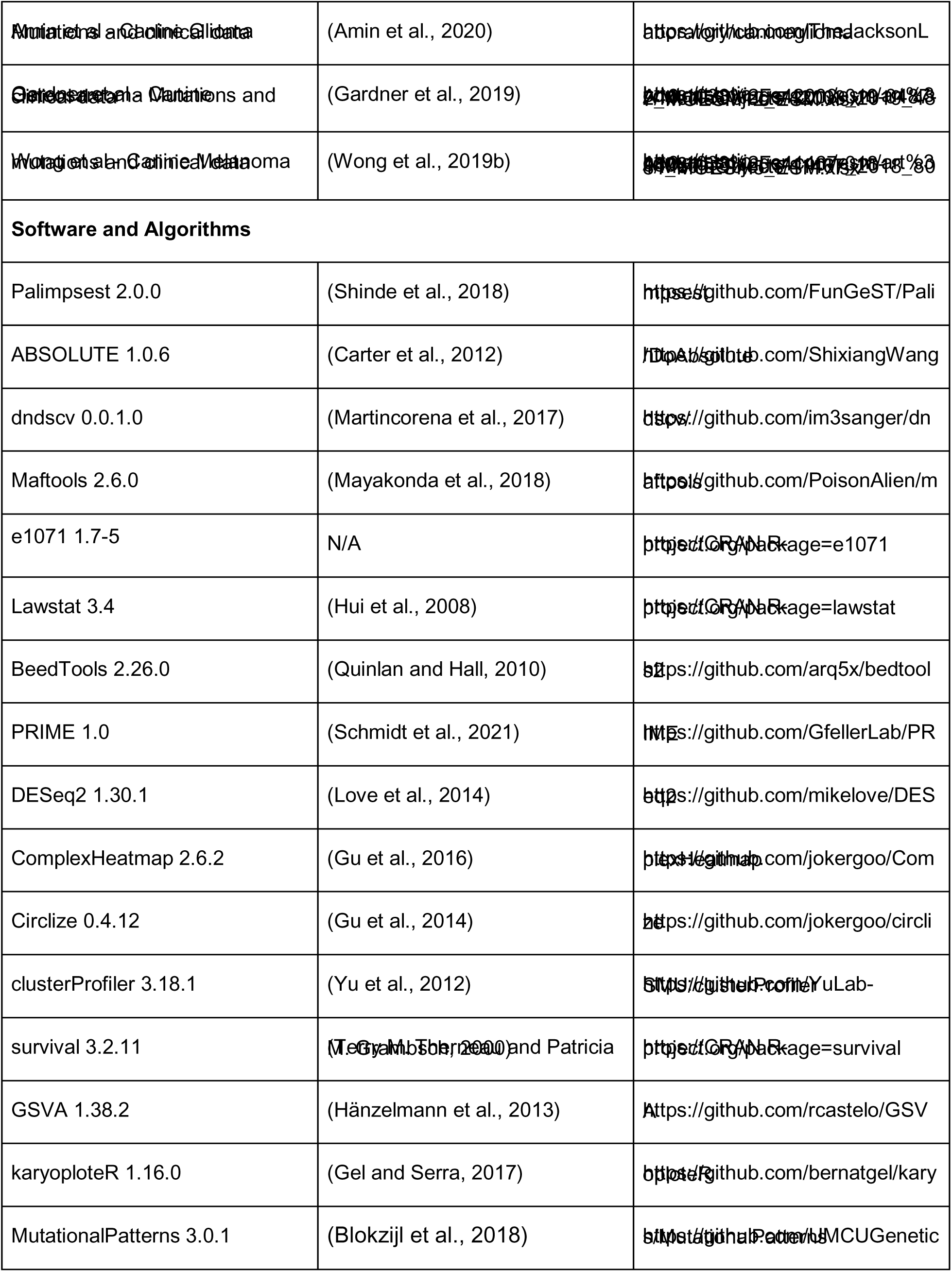
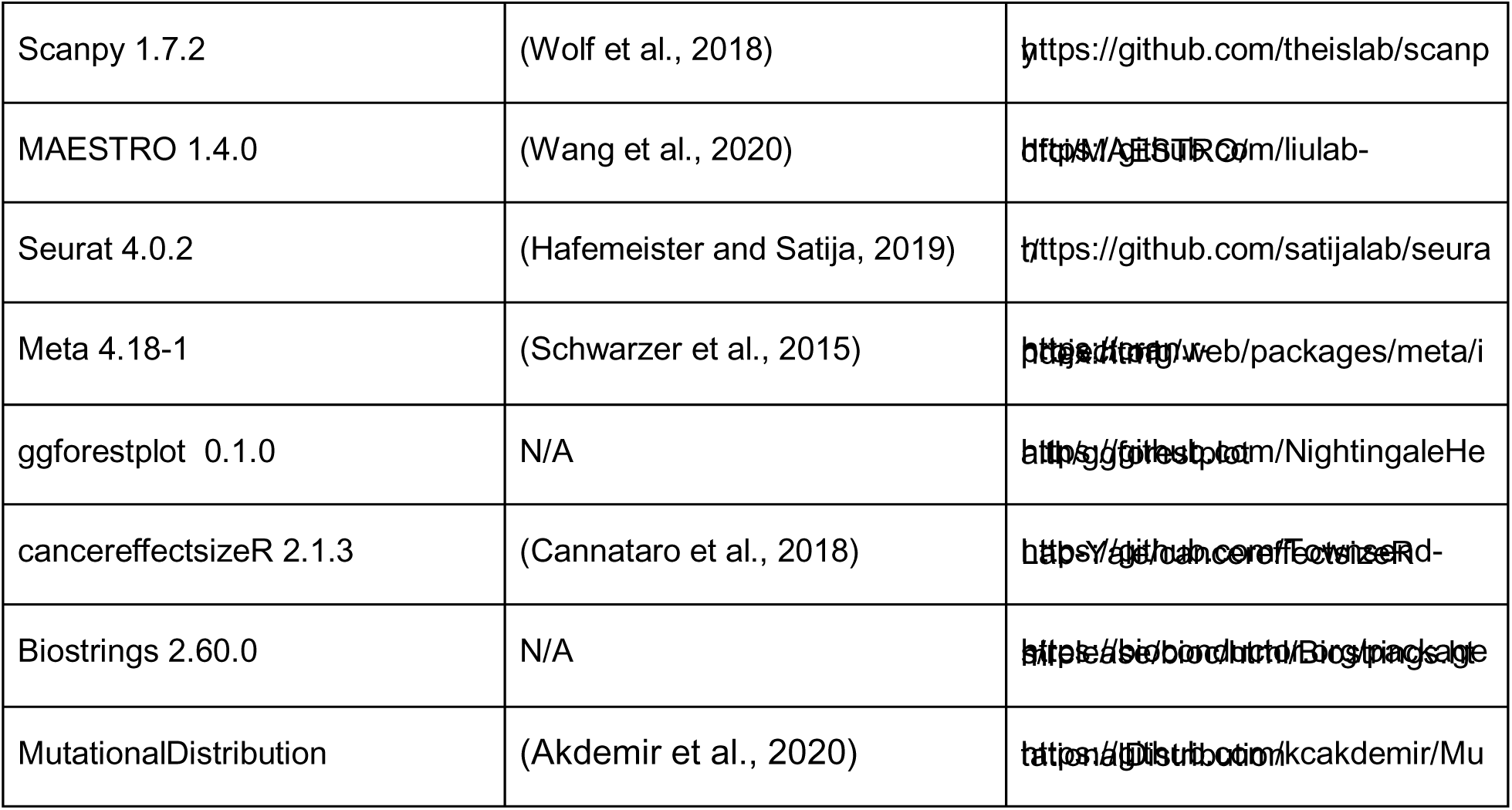

### Data and Code Availability

Our findings are supported by data that are available from public online repositories, or data that are publicly available upon request from the data provider. The code generated during this study is available at https://github.com/iS4i4S/Landscape-AICDA-mutations. The supplementary material is also available at https://data.mendeley.com/drafts/bg9dsdw33y?folder=19693757-9f05-45b5-9c38-87d85f6250ec. All other materials are available upon request from the authors.

### SUBJECT DETAILS

The total cohort at bulk level consisted of 50,631 tumor samples representing more than 80 cancer types (Figure 1A). TCGA information consisted of: mutational data in Mutation Annotation Format (MAF) included 9,264 cancer patients (24 cancer types) and 741 normal samples; RNA-seq data (read counts, n = 9,101); immune data (i.e. cibersort calculated immune populations, n = 8,983), allele-specific integer copy numbers (ABSOLUTE calculated, n = 7,216), available viral counts (n = 5,741) and previously predicted neo-epitopes (n = 2,143; cancer types = 14). The PCAWG data (ICGC) included 2,775 cancer patients along 35 different cancer types with WGS information (SNV and CNV) from which 1,522 had the expression data available. Composite mutations data included 31,353 cancer patients from the MSKCC comprising 41 tumor types by the MSK-IMPACT assay (sizes depending on the date of sequencing comprising 341, 410, and 468 cancer-associated targeted genes) downloaded from CBioPortal for the general maf or their github repository (https://github.com/taylor-lab/composite-mutations/tree/master/data) for the clinical, mutational burden classification, mutational signatures, composite mutation annotation, phasing information and molecular timing (Gorelick et al., 2020). Additionally, hematological cancers cohort (AML, DLBCL, Myelodysplastic Syndromes and other leukemias; n = 3,859) (for The St. Jude Children’s Research Hospital–Washington University Pediatric Cancer Genome Project et al., 2015, 2015; Holmfeldt et al., 2013, 2013; Landau et al., 2013, 2015, 2015; Lohr et al., 2014, 2014; Nangalia et al., 2013, 2013; Papaemmanuil et al., 2016; Puente et al., 2015, 2015; Quesada et al., 2012, 2012; Reddy et al., 2017; the St. Jude Children’s Research Hospital–Washington University Pediatric Cancer Genome Project et al., 2016, 2016; Tyner et al., 2018, 2018, 2018; Welch et al., 2016, 2016; Yoshida et al., 2011, 2011) and pediatric cancers cohort (20 tumor types; n = 1,051) (Gröbner et al., 2018) were obtained from CBioPortal some were part of the Therapeutically Applicable Research to Generate Effective Treatments (TARGET) initiative (phs000467, phs000471 and phs000465). ICI cohort consisted of 2,261 samples coming from: MSKCC-IMPACT dataset (n = 1,472; 11 tumor types), Pender et al. cohort (n = 98, 19 tumor types), Miao et al. cohort (n = 249, four six tumor types), Liu et al. (n = 144, melanoma), Hugo et al (n = 37, melanoma) and Braun et al (n = 261; ccRCC)(Braun et al., 2020a; Hugo et al., 2016b; Liu et al., 2019; Miao et al., 2018b; Pender et al., 2021b; Samstein et al., 2019b). Riaz et al. melanoma cohort consisted of 68 patients treated with Nivolumab (anti-PD-1) from which 35 had previously progressed on Ipilimumab (anti-CTLA-4) treatment from which data was obtained prior treatment (pre) or 4 weeks after initiation of Nivo (on). Data consisted of WES, neo-epitopes (n_pre_ = 68; n_on_ = 41) and RNA-seq (n_pre_ = 45; n_on_ = 41) (Riaz et al., 2017).

The single cell resolution cohort comprised a total of 2,375,926 cells representing 18 tumor types and the human, mouse and PBMC normal cell atlas (Figure 7). The human cell landscape study consisted of 599,926 cells covering 60 tissue types and 63 cell subtypes (Han et al., 2020a); meanwhile the human PBMC atlas included 68,579 cells through 11 cell subtypes (Zheng et al., 2017b). The mouse cell atlas study included 333,778 cells along 47 tissues (Han et al., 2018). Addedly, the oncogenic single cell studies (n = 41; 40 human and one mouse), that included a total of 1,374,222 cells across 18 tumor types with different treatment and sorting conditions (see Key Resources Table and Figure 7F), were obtained mainly from TISCH portal (Sun et al., 2021).

The canine cohort consisted of a total 187 samples representing 3 tumor types including glioma (n = 81), osteosarcoma (n = 35) and melanoma (n = 1) (Amin et al., 2020; Gardner et al., 2019; Wong et al., 2019b).

## METHOD DETAILS

### Tracking *AICDA*-related mutations

We developed a code to detect *AICDA*-related mutations over *wrCy/rGyw* (+/-strand, where "W" stands to either adenine or thymine, "R" to purine and "Y" to pyrimidine) motifs, giving a total of 8 motifs per strand (positive strand = AACC, AACT, AGCC, AGCT, TACC, TACT, TGCC, TGCT; negative strand = TTGG, TTGA, TCGG, TCGA, ATGG, ATGA, ACGG, ACGA), and its enrichment around a 60 bp flanking sequences (the code allows other bp windows for the user). The enrichment strength over the wrCy/rGyw motifs was calculated as:

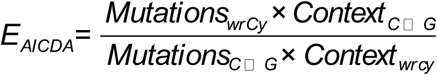

Then we applied a Fisher’s exact test to evaluate the over-representation of AID mutations in each samples by comparing the ratio of substitutions in and out of the AID prefered motifs to the ratio of all cytosines and guanines occurring within the provided genome window around the mutation (60 bp by default), similar to what was previously described for APOBEC (Roberts et al., 2013).

The code was developed under R, takes a maf (mutation annotation format) object as input and outputs a S3 class object containing: i) a matrix of the 768 possible tetranucleotide substitutions across the samples; ii) a data table with all the needed values for enrichment calculation, the enrichment score, Fisher exact-test p-value and fdr for enrichment, fraction of AID mutations, among others; and iii) a maf like data table, with the same format as the input, containing only the attributed AID mutations. Finally, mutations were tagged as AICDA or not AICDA if overlapping or not with the mutations found in the output maf after applying the function.

### *AICDA* motifs distribution across the genome

The *AICDA* motifs, i.e. *WRCY* motifs W=adenine or thymine, R=purine, C=cytosine, Y=pyrimidine, that is: AACC, AGCC, AACT, AGCT, TACC, TGCC, TACT and TGCT. We used a R script to find these patterns in the GRCh38 genome using the Biostrings package v2.60.0. In addition, we also calculated the number of these *AICDA* motifs in 20kb binned windows throughout the genome using bedtools, adjusting by the chromosome size.

### Attributing mutations to mutagenic processes

Mutations previously tagged as not AICDA were subjected to signature attribution to 46 (we excluded signatures SBS: 27, 39, 43, 45-60 since they are attributed to sequencing artefacts) of the 65 COSMIC mutational signatures (v3.0) using *Palimpsest* package with default parameters (Alexandrov et al., 2013, 2020; Shinde et al., 2018). To avoid over-fitting, signatures not contributing with at least one mutation within 50% of the samples per tumor type (Median-SBS_n_^TumorX^ < 1) were removed and mutations were re-fitted using the remaining signatures. Furthermore, signatures proportions per sample were re-calculated adding the number of previously identified AID mutations to the signature data. Signatures were not calculated from the MSKCC-Composites and ICI cohorts because they were already available or not used.

### Clonality analysis

Clonal and subclonal categorization of mutations was done only on the TCGA and PCAWG (only somatic chromosomes) cohorts from which allele-specific integer copy numbers, ploidy and purity estimates were available (n = 7,216 and n = 2,707, respectively). As previously described, for each mutation the cancer cell fraction (CCF) along with the 95% CI (binomial distribution) was determined through the variant allele fraction, tumor purity and local copy numbers, then we assigned a mutation (using *Palimpsest* package) as subclonal if the upper boundary of the 95% CI CCF value was inferior to 0.95, or clonal otherwise (Shinde et al., 2018).

### Attributing viral counts to TCGA samples

As previously described, we assigned samples as being infected by EBV, HBV, HCV, HPV, Kaposi virus or polyomavirus if the viral reads per million reads mapped to the human genome exceeded the maximum observed from normal samples (GTEx cohort), this based on the idea of low levels representing previously exposed leukocytes in the normal tissue sample (Rooney et al., 2015).

### Replication timing

Replication timing measurements of 11 different cell lines (SK-N-SH = neuroblastoma; MCF-7 = mammary gland adenocarcinoma; BJ = skin fibroblast; NHEK = epidermal keratinocytes; HepG2 = liver carcinoma; IMR90 = fetal lung fibroblasts; K562 = leukemia; HeLa-S3 = cervical carcinoma; GM12878 = lymphoblastoid; HUVEC = umbilical vein endothelial cells; BG02ES = embryonic stem cell) were downloaded as wavelet-smoothed signal and then used to compute the median Repliseq signal within 1 Mb or 25 Kb (for TADS) windows which resulted in values from 0-100 indicating late to earlier replication times. We further proceeded to divide the genome in 1 Mb regions (or 25 Kb) by first masking out all regions in the genome that requires 36-mer to be unique in the genome even after allowing for two differing nucleotides and also the Duke and DAC blacklisted regions (highly possible anomalous signals) using BEDTools (Quinlan and Hall, 2010), this led to 3,053 1 Mb windows. Then we calculated the mean number of mutations attributed to AICDA or APOBEC (COSMIC SBS2 and SBS13), per tumor type, as the mutation number within each bin divided by the number of samples within the corresponding tumor type, then we coupled the corresponding repliseq data (as decile distributed) for each bin. For the TCGA data, the mutation density was adjusted by multiplying by the effective exonic length (EEL) fraction (EEL divided by the window width).

The distribution of mutations was divided in 70 bins for plotting, skewness was calculated by Pearson’s moment coefficient of skewness (e1071 R package) and symmetry was tested using a robust estimate of the standard deviation combined with 1,000 bootstrapping to estimate sampling distribution, as proposed previously (Miao et al., 2006) (implemented in lawstat R package) in which left or right hypothesis was set based on previously calculated skewness. Late/early replication was considered enriched if the p-value was inferior to 0.05.

Samples were classified as having microsatellite instability (MSI) or as microsatellite stable (MSS) according to the PCAWG Technical Working Group previous assignment. Only tumor types having more than 20 MSI samples were used for comparing the effect of defective MMR machinery on AICDA or APOBEC induced mutations (1,527 samples, 9 tumor types).

### Mutation distribution around TAD boundaries

To compare the mutational load distribution generated by different mutational signatures, we calculated the ratio of mutation burden in transcriptionally inactive domains versus transcriptionally active domains. First, for each sample, we binned the mutations in 25-kb nonoverlapping windows along the genome. Next, we calculated the sum of mutations at inactive and active domains and normalized this value by the length of active and inactive domains, as previously described (Akdemir et al., 2020).

### Clonal spectral changes, signature changes and clonal gene enrichment

First we assessed which samples had measurable mutation spectral changes between clonal mutations and subclonal mutations by assuming that the trinucleotide single base substitution spectra follow a multinomial distribution and comparing the early (clonal) versus late (subclonal) spectra, as previously described (PCAWG Evolution & Heterogeneity Working Group et al., 2020). From the resulting samples, we measured the fold change of relative proportion signature activities (subclonal to clonal) per sample as:

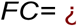

Where the relative activity is the quotient from the number of mutations due to a signature and the total number of mutations, per timing (subclonal or clonal).

Clonality enrichment per gene (g) was calculated by comparing the odds ratio, assuming an hypergeometric distribution, of the clonal/subclonal (c/sc) mutations observed (n_o_) to those expected by chance (n_c_):

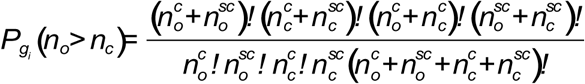

Negative binomial regression was used for estimating n_c_ per gene and as clonal 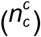 or subclonal 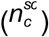 by modeling observed number of clonal/subclonal mutations adjusted for coding sequence length (l), GC content percentage (g, Biomart GRCh37), replication time (r), chromatin state (h) (Lawrence et al., 2013), the average total DNA copy number of the gene across its mutated samples (tcn), the cancer cell fraction (ccf) and sample purity (pu). Additionally, an offset term was added to the model that represents the log-number of tumor samples (n_s_) harboring mutations in the gene of interest. The final model can be represented as:

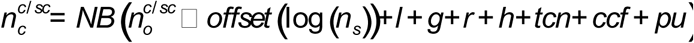

Genes having and OR > 1 were considered more clonal, or more subclonal if inferior to 1 and superior to 0. Clonal enrichment was achieved if having a FDR corrected p-value (Fisher one-sided test) less than 0.01.

### Attributing mutations as driver genes

Positively-selected genes per tumor type within each cohort, excluding MSKCC-Composites from which we used the already curated list from the original article (Gorelick et al., 2020), were obtained by calculating dN/dS likelihood ratios (*dNdScv* package) through negative binomial regression modeling of the background mutation rate of each gene using distinct genomic covariates including variation in mutation density across genes, context-dependent substitutions (mutational signatures), transcriptional strand bias, chromatin state, expression and replication time. Additionally, we removed ultra-hypermutator samples and extremely mutated genes per sample, to avoid loss of sensitivity. Genes were considered as drivers if having q-values < 0.01 (Benjamini-Hochberg’s multiple testing correction of p-values) (Martincorena et al., 2017). In addition, in the ICGC cohort, we also used the selection intensity of every particular mutation related with APOBEC or AICDA signatures by deconvolution of prevalence by mutation rates for recurrent amino acid mutations within three oncoproteins caused by single-nucleotide changes using cancereffectsizeR v2.1.3 package. The observed substitution rates were divided by the expected substitution rates in the absence of selection. The expected substitution rates in the absence of selection were calculated as the average per-site synonymous mutation rate of the gene, normalized for the average weight of trinucleotide mutational signature burden for that signature. The quotient of observed to expected numbers of substitutions was the selection intensity, as previously described (Cannataro et al., 2018).

### Gene and residue-specific composite mutation enrichment testing

Main code for analysis was obtained from the original article and then subjected to minor modifications for identification of *AICDA*-related mutation’s contribution (Gorelick et al., 2020). In brief, the number of samples harboring a composite mutation in permutation i (n_i_) was obtained by permuting 100,00 times the z component of a matrix of m x z (m = total number of nonsynonymous somatic mutations; z = mutation id as mutation in x gene occuring in a x sample). A p-value was calculated as the fraction of permutations satisfying n_i_ ≥ n^pos^.

Enrichment of AID composites per gene was assessed using a binomial test to evaluate, for each gene, that the proportion of observed AID composite mutated samples differs from the proportion of predicted AID composite mutated samples arising by chance (n_c_). Negative binomial regression was used for estimating n_c_ per gene by modeling observed number of AID composite-mutant samples adjusted for multiple genomic covariates like coding sequence length (l), GC content percentage (g, Biomart GRCh37), replication time (r), chromatin state (h) (Lawrence et al., 2013), MSK-IMPACT targeted assay version (i) and the average total DNA copy number of the gene across its mutated samples (t). Additionally, an offset term was added to the model that represents the log-number of tumor samples harboring mutations in the gene of interest. Enrichment at individual mutant residues arising as AID composite mutations were assessed using a right-sided Fisher’s exact test comparing if it arose significantly more frequently than all other mutant residues within the same gene. Residues and genes were considered significant if having a FDR corrected p-value less than 0.01.

### Mutations distribution within R-loops and transcription correlation

Genomic regions coordinates file with GC skew enrichment, which are associated with R-loop formation, downloaded from a previous study as GRCh38 assembly, was transformed to GRCh19 assembly using USCS genome browser liftover function (Ginno et al., 2012; Kent et al., 2002). This gave rise to 16,223 regions (median 626 bp) from which only G skewed regions (8,059; associated with R-loops formation during transcription) were used to find overlaps with the genomic position of each mutation and to label them as “in R-loops” or “out R-loops”.

Transcription correlations with *AICDA*-related mutations were performed by Spearman correlation method. For each tumor type, we correlated the number of *AICDA*-related mutations occurring in gene i (AID_Muts_gi_) to the expression of the same gene i (Exp_gi_). The AID_Muts_gi_ were considered associated with the transcription process in a specific tumor type if the adjusted p-value was inferior to 0.05 and the spearman Rho value superior to 0. Functional enrichment analysis of the associated genes was performed using the DAVID database (Huang et al., 2009).

### Neoepitope analysis

TCGA neoepitope list and HLA haplotypes were retrieved from previous studies, in bref 4-digit HLA type for each sample was inferred using POLYSOLVER and neoepitopes were predicted using NetMHCpan (v2.4) (only for MHC class I) as novel 9-10mers that resulted from mutations in expressed genes (>10 TPM) and affinity < 500 nM (Rooney et al., 2015; Shao et al., 2020). We coupled this information with the data resulting from our analysis to obtain neoepitopes data (n = 3,370 patients) with clonality, signature origin, expression, PolyPhen/SIFT effect on coding protein, among others. We furthered filtered for >1 FPKM expression on the genes originating the neopeptide and excluded patients with: I) incomplete HLA information; II) microsatellite instability (retrieved from Ding et al., 2018) (Ding et al., 2018); III) altered antigen presentation related genes *HLA-A, HLA-B, HLA-C, CIITA, IRF1, PSME1, PSME2, PSME3, ERAP1, ERAP2, HSPA, HSPC, TAP1, TAP2, TAPBP, CALR, CNX, PDIA3 and B2M* (HLA enhanceosome, peptide generation, chaperones and the MHC complex) which was based if presence of PolyPhen/SIFT damaging predictions (‘‘probably_damaging’’/‘‘possibly_damaging’’ or ‘‘deleterious’’/‘‘deleterious_low_confidence’’, respectively) or of copy number loss. These filters left 2,143 patients that were considered for Figure 6A and 6B and Supplementary Figure 22. To account for immunogenicity based on prediction of neopeptide TCR recognition and neopeptide HLA binding, PRIME software was run with default parameters. Neoepitopes were classified as “Immunogenic” if having a PRIME %rank score (the fraction of random 700,000 8- to 14-mers that would have a score higher than the peptide provided in input) lower or equal to 0.5% for the corresponding HLA haplotype of the patient where the neopeptide occurred, or as “Non-Immunogenic” otherwise (Schmidt et al., 2021). For simplicity some cosmic signatures were grouped together as: MMR (SBS6, SBS15, SBS20, SBS21, SBS 26 and SBS44); Smoking-associated (SBS4, SBS18, SBS24 and SBS29); POLE (SBS10a, SBS10b and SBS14) and APOBEC (SBS2 and SBS13). This data combined with the clonality of the mutation giving rise to a specific neopeptide was used to classify them as immunogenic clonal neoepitope (ICN); samples were further classified as “Presence” if having at least one ICN due to a specific mutational signature or “Absence” otherwise. TCGA RNA counts were retrieved from GEO GSE62944 and normalized using the variance-stabilizing transformation (VST) function from DESeq2 (Love et al., 2014).

For the advanced melanoma anti-PD-1 treated cohort (Riaz et al., 2017) mutational signatures, clonality, immunogenicity and RNA counts normalization was assessed as described above. Samples were classified as having high ICN AID load if their load was superior to the cohort’s median. Furthermore, differential gene expression analysis (Ipi-Naive samples only) was executed with DESeq2 comparing high ICN AID load versus low within pre-therapy samples only (Figure 6F; Supplementary Figure 23) or comparing pre-therapy to on-therapy adjusting for ICN AICDA load groups (high or low) to identify expression changes on-therapy (design = ∼CIN_AID_binary+PreOn). For each comparison, DEGs (adjusted p-value <0.2) were used as input for hierarchical clustering (Euclidean distance followed by complete-linkage agglomeration algorithm) to obtain gene clusters used for enrichment analysis (Figure 6 E,F). GSEA analysis was performed using the GO database through R package clusterProfiler (Yu et al., 2012) or DAVID database (Huang et al., 2009) applying Bonferroni correction (q-value <0.05).

### Analysis of single cell RNA sequencing data

We downloaded the expression matrix of the raw count, transcript per million (TPM) or Fragments Per Kilobase of transcript per Million mapped reads (FPKM), if available for each dataset. We collected sample information from databases or the original studies, such as the patient ID, tissue origin, treatment condition and response groups.

For the cancer related datasets, we used a standardized analysis workflow based on MAESTRO v1.4.0 (Wang et al., 2020) for processing all the collected datasets, including quality control, batch effect removal, cell clustering, differential expression analysis, cell-type annotation, malignant cell classification and gene set enrichment analysis that internally uses the Seurat package (Hafemeister and Satija, 2019). The raw count, TPM or FPKM table was used as input for the standardized workflow. The quality of cells was determined by two metrics: the number of total counts (UMI) per cell (library size) and the number of detected genes per cell. Low-quality cells were filtered out if the library size was <1000, or the number of detected genes was <300. For each cancer related dataset, the MAESTRO workflow identified the top 2000 variable features and employed principal component analysis (PCA) for dimension reduction, k-nearest neighbors algorithm (k-NN), and Louvain algorithm for identifying clusters (Stuart et al., 2019; Xu and Su, 2015). To better capture the cellular difference and variabilities for datasets with different cell numbers, we adjusted the number of principal components and the resolution for graph-based clustering, which were both increased with the cell number. The uniform manifold approximation and projection (UMAP) were utilized to reduce the dimension further and visualize the clustering results (Becht et al., 2019). We used the cell annotation provided by TISCH database (Sun et al., 2021b) and when the study was not included in TISCH we followed the same methodology described in TISCH to homogeneously annotate the cells throughout the different cancer related studies.

Similarly, for the non-cancer studies: the Mouse Cell Atlas (MCA) (Han et al., 2018b), the Human Cell Landscape (HCL) (Han et al., 2020b) and the analysis of 68k fresh peripheral blood mononuclear cells (PBMCs) (Zheng et al., 2017c) we used Scanpy v1.7.2 (Wolf et al., 2018) with the aforementioned filters and dimension reduction methods. We used the cell annotation, the tissue of origin and the stage (i.e. adult, fetal, etc), when available, provided by the authors of each study. The hierarchical clustering by tissue, cell type or stage of differentiation was performed using the *matrixplot* function in Scanpy using default parameters.

The cell cycle analysis was estimated using the *CellCycleScoring* function from the Seurat package using a list of genes related to the cell cycle (Tirosh 2016). The algorithm calculates the difference of mean expression of the given list and the mean expression of reference genes. To build the reference, the function randomly chooses a list of genes matching the distribution of the expression of the given list. Cell cycle scoring adds three slots in data, a score for S phase, a score for G2M or for G1 phase. Similarly, in the datasets analyzed with Scanpy, we used *sc.tl.score_cell_cycle_genes* function using the same gene list provided by Seurat.

### Statistical analyses and figures

All statistical analyses were performed using the R statistical programming environment (version 4.0). Figures were generated using either base R or the ggplot2 library. Differences in proportions were calculated from Fisher’s exact test or two-sample Z-tests. Error bars indicate the 95% binomial CIs calculated using the Pearson-Klopper method. Kruskal-Wallis test was used to test for a difference in distribution between three or more independent groups, and Mann Whitney U test was used for differences in distributions between two population groups, unless otherwise noted. Spearman correlations were calculated by the cor.test function in R. P-values were corrected for multiple comparisons using the Benjamini-Hochberg method when applicable. For heatmap representation (ComplexHeatmap or circlize R packages)(Gu et al., 2016), VST gene expression values were first quantile normalized and log2 transformed and then converted to Z-scores. Overall survival analysis to ICI was assessed using log-rank Kaplan-Meier curves and univariate/multivariate Cox proportional hazards regression modeling. We have assessed several Cox proportional model for every study (i.e. analyzing the deciles, from 10th to 90th, of the fraction of AID induced mutations in every included study, unadjusted, using the median of the fraction of AID mutations, and also these models were adjusted by TMB >= 10mut/Mb). To combine the different survival models, we used a random-effects model with the meta v4.18-1 package (Schwarzer et al., 2015), using log hazard ratio and standard errors of each model per study. The inverse variance method was used for pooling. The random-effects estimate was based on the DerSimonian-Laird method (DerSimonian and Laird, 2015). The meta-analysis results were represented in a forestplot using the *forestplot* function of the ggforestplot v0.1.0 package.

---

**Supplementary Figure 1. AID-related mutations distribution across cohorts**

Summary of the frequency of the different COMIC SBS signatures, including the signature related with AID within the TCGA (Panel A), Pediatric (Panel B) and hematological (Panel C) datasets. Panel D, distribution of the SBS signatures contribution in the different cancer types of the TCGA dataset per sample where AICDA signature is represented in black with a smooth regression line in red (related to Panel A).

**Supplementary Figure 2. Mutational signatures distribution and AICDA/APOBEC mutations frequency across cohorts**

Distribution of the SBS signatures contribution in the different cancer types of the pediatric (Panel A) and hematological (Panel B) dataset per sample where AICDA signature is represented in black with a smooth regression line in red (related to Supplementary Figure 1). Frequency of AID and APOBEC-related mutations in the TCGA (Panel C), MSKCC (Panel D), Pediatric (Panel E) and Hematological (Panel F) cohorts.

**Supplementary Figure 3. AICDA mutations genomic distribution TCGA (Tumors ACC, BLCA, BRCA, CESC, COAD and DLBCL)**

Rainfall plots of the AICDA mutations’ distribution across chromosomes on TCGA samples by tumor types as a function of logarithmic (10 scale) genomic distance. Black points represent C to G mutations and red points C to T mutations. Top barplot shows the sum of mutations (density) across chromosomes.

**Supplementary Figure 4. AICDA mutations genomic distribution TCGA (Tumors GBM, HNSC, KICH, KIRC, KIRP and LAML)**

Rainfall plots of the AICDA mutations’ distribution across chromosomes on TCGA samples by tumor types as a function of logarithmic (10 scale) genomic distance. Black points represent C to G mutations and red points C to T mutations. Top barplot shows the sum of mutations (density) across chromosomes.

**Supplementary Figure 5. AICDA mutations genomic distribution TCGA (Tumors LGG, LIHC, LUAD, LUSC, OV and PRAD)**

Rainfall plots of the AICDA mutations’ distribution across chromosomes on TCGA samples by tumor types as a function of logarithmic (10 scale) genomic distance. Black points represent C to G mutations and red points C to T mutations. Top barplot shows the sum of mutations (density) across chromosomes.

**Supplementary Figure 6. AICDA mutations genomic distribution TCGA (Tumors READ, SKCM, STAD, THCA, UCEC and UCS)**

Rainfall plots of the AICDA mutations’ distribution across chromosomes on TCGA samples by tumor types as a function of logarithmic (10 scale) genomic distance. Black points represent C to G mutations and red points C to T mutations. Top barplot shows the sum of mutations (density) across chromosomes.

**Supplementary Figure 7. AICDA motifs/mutations genomic distribution in normal genome or DLBCL tumors**

Genomic distribution of AICDA related motifs across chromosomes as 20 kb window raw counts (Panel A) or counts adjusted by chromosome length (Panel B). Chromosome 2 was used as the reference group for Wilcoxon-test (FDR adjusted p-values). Panel C-F, rainfall plots of the AICDA mutations’ distribution in DLBCL tumors (TCGA cohort) on chromosomes 14, 22, 2 and 6, respectively.

**Supplementary Figure 8. AICDA mutations contribution on driver genes**

Mutational signatures’ contribution (COSMIC and AICDA signatures) to driver genes on the ICGC (Panel A), TCGA (Panel B), Pediatric (Panel C) and Hematological (Panel D) cohorts across tumor types where black bars represent AICDA signature.

**Supplementary Figure 9. Selection intensity of AICDA mutations**

AICDA mutations produce higher selection intensity on driver genes on minor hotspot residues but there is a higher number of affected genes/residues than the ones generated by the APOBEC related signatures.

**Supplementary Figure 10. AICDA expression/mutations correlations with other features (TCGA cohort)**

Spearman correlations of AICDA expression with mutations (Panel A) and age (Panel C) or AICDA mutations with TMB (Panel B). Panel D, cumulative distribution of age according to AICDA or APOBEC signatures. Panel E, age distribution comparison according to gender of AICDA/APOBEC mutational signatures (Wilcoxon-test).

**Supplementary Figure 11. AID-mutations interplay with replication across tumor types (ICGC cohort)**

Distribution of late to early replication time according to AICDA (Panel A), and APOBEC signatures SBS2 (Panel B) and SBS13 (Panel C), related to Figure 3A.

**Supplementary Figure 12. APOBEC mutations interplay with TADs**

Average profile of SBS2 (Panel A) and SBS13 (Panel B) somatic mutations accumulation in 2,775 cancer samples and replication timing across 500 kb of TAD boundaries delineating active to inactive domains. Related to Figure 3 B-D.

**Supplementary Figure 13. AID-mutations interplay with R-loops, transcription and clonality**

Panels A and B, average profiles of SBS2 or SBS13 induced mutations accumulation in 2,775 cancer samples across 500 kb of TSS for negative strand genes (top) or positive strand genes (bottom). Panel C, percentage of clonal/subclonal mutations attributed to AICDA or APOBEC mutational signatures (Two sided Fisher’s exact test). Panel D, fold change of signature activities (subclonal to clonal) across tumor types for AICDA, APOBEC (SBS2 and SBS13) and SBS9 mutational signatures for samples with measurables changes in their mutation spectra (n = 404) according to early mutations (top) or late mutations (bottom) where box plots indicate the first and third quartiles of the distribution, with the median shown in the centre and whiskers covering data within 1.5x the IQR from the box. Related to Figure 3E-G.

**Supplementary Figure 14. AICDA mutations role in composite mutations**

Panel A, temporal order of acquisition of global composite mutations by cancer gene function using clonality and allelic configuration (Two-sided binomial test; Related to Figure 4D). Panel B, mutational signatures contribution to composite mutations globally (left) and per driver gene (right).

**Supplementary Figure 15The impact of AID/APOBEC mutations on ICI response**

Meta-analysis of the survival impact of the fraction of AICDA mutations in different studies. Panel A, effect using all the deciles of the fractions of AICDA mutations at univariate level, the overall impact of AICDA with a better OS is present independently of the cut-off (relative to Figure 5B). Panels B and C, assessment of the prognostic value of the fraction of AICDA mutations in the IMPACT study. Forest plot of a Cox model of either the global impact, after adjustment by TMB (≥10 mut/Mb), median APOBEC mutations, age and gender (B) or the impact by tumor subtype (C), relative to Figures 5C and 5D. Panels D-F, correlation between the fraction of AICDA mutations and the fraction of APOBEC mutations in the IMPACT (D), ICGC (E) or TCGA (F) datasets.

**Supplementary Figure 16.** Results from Miao et al study Panel A) forest plot showing the overall impact estimated using univariate Cox proportional hazard ratio model (across all samples) of TBM and the fractions AICDA mutations on overall survival (top of the figure) and per every type of cancer included in this study. Panel B) Forest plot with the results of the multivariate Cox proportional hazard ratio model of AICDA related mutations with OS, after adjusting by TMB, age and sex.

**Supplementary Figure 17.** Results from Liu et al study. Panel A) Kaplan-Meier plot showing a better overall survival in patients with higher fraction of AICDA mutations (according to the median). Panel B) Distribution using boxplots of the fraction of AICDA mutations according to the best response under ICI treatment, PD (progression disease), PR/CR (partial response/complete response) and SD/MR (stable disease/mixed response). Panel C) The distribution using boxplots of the fraction of AICDA mutations according to the localization of the melanoma. Panel D) Forestplot of the Cox proportional hazards ratio multivariate model adjusting the potential prognostic impact on OS of the fraction of AICDA mutations (according to the median) with tumor purity and gender.

**Supplementary Figure 18.** Results from Pender et al. study. Panel A) Forestplot of the Cox proportional hazard ratio multivariate model to assess the prognostic impact on OS of the fraction of AICDA mutations (according to the median), adjusted by TMB, age and gender. Panel B) The distribution of the fraction of AICDA mutations according to the clinical benefit of using ICI, defined using NCB (no clinical benefit) and DCB (durable clinical benefit).

**Supplementary Figure 19.** Results from Hugo et al. study. Panel A) Kaplan-Meier plot comparing the OS according to the fraction of AICDA mutations (cut-off median). Panel B) Forestplot of the Cox proportional hazard ratio multivariate model to assess the prognostic value of the fraction of AICDA mutations on OS adjusting by TMB (cut-off ≥ 10).

**Supplementary Figure 20.** Results from the Braun et al. study. Panel A) Kaplan-Meier plot comparing the OS according to the fraction of AICDA mutations (cut-off media). Panel B) Forestplot of the Cox proportional hazard ratio multivariate model to assess the prognostic value of the fraction of AICDA mutations on OS adjusting by gender, age (cut-off median) and PBRM1 mutations.

**Supplementary Figure 21. Landscape of AID-related neoepitopes (TCGA cohort)**

Panel A, clonal versus subclonal immunogenic neoepitopes comparison due to different mutational signatures across tumor types (Wilcoxon test). Panel B, comparison of the number of clonal immunogenic neoepitopes produced by the different mutational signatures across tumor types (Wilcoxon test).

**Supplementary Figure 22. AID-related neoepitopes role in ICI response (Riaz et al cohort)**

Panel A, OS prediction within Ipi-Naive patients using mutational load (top left) or within Ipi-Progressive patients using UV mutational load (top right), AICDA mutational load (bottom left) or global mutational load (bottom right). Panel B, heatmap and hierarchical clustering of DEGs between high and low ICN AICDA load within pre-therapy Ipi-Naive patients. Panel C and D, boxplots comparison showing increased expression trend of either inhibitory immune checkpoint molecules or cytolytic activity, respectively (Wilcoxon-test). Relative to Figure 6.

